# The antagonism of sex chromosomes and sex hormones dictates outcomes in Th17 cell-mediated central nervous system autoimmunity

**DOI:** 10.64898/2026.11.09.765425

**Authors:** Muhammad Umair, Joanie Baillargeon, Charles Joly-Beauparlant, Mohamed Reda Fazazi, Alexia Falle, Krish Pandey, Pascal Dufour, Rene Quevedo, Ebenezer Asiedu, Irshad Akbar, Prenitha Mercy Ignatius Arokia Doss, Vahid Safdari, Camille Lavoie-Ouellet, Patrick Caron, Ana C. Anderson, Chantal Guillemette, Clémence Belleannée, Connie M. Krawczyk, Tracy L. McGaha, Arnaud Droit, Manu Rangachari

**Affiliations:** axe Neurosciences, Centre de recherche du CHU de Québec–Université Laval, Pavillon CHUL, 2705 boulevard Laurier, Quebec City, QC G1V 4G2, Canada; axe Endocrinologie et Néphrologie, Centre de recherche du CHU de Québec–Université Laval, Pavillon CHUL, 2705 boulevard Laurier, Quebec City, QC G1V 4G2, Canada; Tumor Immunotherapy Program, Princess Margaret Cancer Centre, Toronto, ON M5G 2M9, Canada; Department of Immunology, University of Toronto, Toronto, ON M5S 1A8, Canada; Department of Metabolism and Nutritional Programming, Van Andel Institute, Grand Rapids MI, USA; axe Reproduction, santé de la mère et de l’enfant, Centre de recherche du CHU de Québec–Université Laval, Pavillon CHUL, 2705 boulevard Laurier, Quebec City, QC G1V 4G2, Canada; Faculty of Pharmacy, Université Laval, 1050 ave de la Médecine, Quebec City, QC, Canada; Gene Lay Institute of Immunology and Inflammation, Harvard Medical School and Mass General Brigham, Boston, MA 02115, USA; Faculty of Medicine, Université Laval, 1050 ave de la Médecine, Quebec City, QC, Canada

**Keywords:** biological sex, sex chromosomes, androgen receptor, Four Core Genotypes (FCG), multiple sclerosis, experimental autoimmune encephalomyelitis, T helper 17 (Th17), Wnt, epigenetic inheritance

## Abstract

Biological sex traits are shaped by the combined effects of both sex hormones and chromosomes. How these factors interact in sex-related autoimmune diseases, such as multiple sclerosis (MS), has been a subject of intense recent interest. Here, using the Four Core Genotype (FCG) mouse model, in which these two variables can be disentangled, we describe an antagonism between sex hormones and chromosomes that potently influences outcomes in T helper 17 (Th17)-driven progressive central nervous system (CNS) autoimmunity. Male gonadal hormones dampen disease severity, irrespective of sex chromosome complement, by promoting a Wnt/β-catenin signaling program associated with a homeostatic, non-pathogenic Th17 phenotype. By contrast, XY chromosomal complement exacerbates Th17 pathogenicity in a recipient sex-dependent manner; we identify the X-linked molecule SUV39H1 as an inhibitor of inflammatory Th17 cell function. Unexpectedly, we found that hormonally and chromosomally female FCG Th17 cells induced far more severe disease than WT female counterparts despite being genetically identical, pointing to a parental imprinting effect in dictating Th17 cell pathogenicity. These latter investigations uncovered Runx2, a transcription factor previously linked to developmental programs, as a novel negative regulator of the pathogenicity of Th17 cells. Together, our findings establish biological sex as a multilayered regulator of Th17 pathogenicity, revealing distinct and sometimes opposing contributions of hormones, chromosomes, and parental inheritance.

## Introduction

Biological sex is a fundamental organizing trait of virtually all eukaryotes. For eutherian mammals, sex is first determined by inheritance of XX (female) vs XY (male) chromosomal complement. Expression of the chromosome Y-encoded testes determining gene, *Sry*, drives a male differentiation program. Thus, in XX organisms that lack *Sry*, a female differentiation program is instead enacted ^1,2^. According to a traditional model of sexual differentiation, the sex hormones (estrogens and progesterone in the case of females, androgens in the case of males) are then critical for enforcing sexual maturity. While this dogma has more recently been called into question^3^, it is nevertheless broadly accepted that sex chromosomes and hormones collaborate to drive dimorphism between biological males and females.

Although biological sex is primarily associated with reproduction, it is increasingly recognized as a fundamental determinant of physiology, influencing multiple organ systems and disease processes. The impact of sex hormones on adult biology is well documented^4^. In contrast, the contribution of sex chromosomes independent of gonadal hormones remains less well understood. A traditional view held that the role of sex chromosomes is primarily to set sexual differentiation in motion during gestation and that they have limited impact in non-reproductive tissues in sexually mature mammals. This belief was supported by the knowledge that the Y chromosome is enriched for genes specific for male fertility, and that X chromosome genes are typically silenced on one copy in females to ensure equivalent dosage with males^5^. However, it is now known that a significant proportion of X-linked genes escape second-allele silencing^6^. Thus, the role of the sex chromosomes in regulating sex differences in adult physiology are becoming increasingly appreciated. Further, while one might expect sex-matched hormones and chromosomes to act synergistically to enforce a given phenotype, it has been observed that they can also work in opposition to one another^7,8^.

Adaptive immune responses are subject to striking sex differences. Females mount more robust responses to viral infection and have increased susceptibility to autoimmune diseases such as rheumatoid arthritis and systemic lupus erythematosus^9^. This has led to a common view that T cell-mediated immunity is more pronounced in females. However, there is increasing evidence that autoimmune T helper 17 (Th17) responses are more potent in males^10,11^. Multiple sclerosis (MS), an autoimmune disease of the brain and spinal cord, shows particularly intriguing sex effects: women develop relapse-onset MS with a 3-fold higher frequency than men, yet men tend to develop more severe symptoms and are at greater risk of developing progressive forms of MS^12^. This latter point is crucial as progressive MS is largely refractory to current therapies^13^.

Here, we used the Four Core Genotypes (FCG) model to separately examine the contribution of sex hormones and chromosomes to this phenotype, using an adoptive transfer model of CNS autoimmunity in which male, but not female, Th17 cells can induce severe chronic disease in recipients^14^. Male gonadal hormones ameliorate Th17-mediated disease via the activity of a Wnt/androgen receptor (AR) axis. By contrast, XY chromosomes exacerbate disease, at least in part due to reduced expression of the X-linked molecule Suv39h1 in male Th17 cells. We additionally find evidence of a transgenerational inheritance effect by which the pathogenicity of Th17 cells from otherwise genetically identical mice is dictated by the genotype of their sires. Together, our data provide evidence of a complex and multilayered regulation of T cell function by biological sex.

## Results

### Male hormones ameliorate Th17 pathogenicity while XY worsens it

Experimental autoimmune encephalomyelitis (EAE) is a well-established mouse model of the immune aspects of MS pathogenesis that can be induced either by active immunization with myelin antigens or by adoptive transfer of myelin antigen-specific T lymphocytes^15^. On the nonobese (NOD) genetic background, a substantial frequency of EAE mice develop a relapsing-to-chronic progressive disease course ^14,16,17^. We recently described an EAE model (schematic in Figure 1A) in which myelin oligodendrocyte glycoprotein (MOG)_[35-55]_-specific 1C6 T cell-receptor-transgenic (TcR-Tg) ^18,19^ NOD-background Th17 cells were adoptively transferred to NOD.*Scid* recipient mice. Curiously, although recipients of female Th17 cells rarely develop progressive disease, male Th17 cells are responsible for a wide divergence of outcomes; approximately 60% of recipients of male Th17 cells develop severe progressive EAE, while the remainder develop much milder disease that is similar in severity to that caused by female cells ^14^.

**Figure 1.**
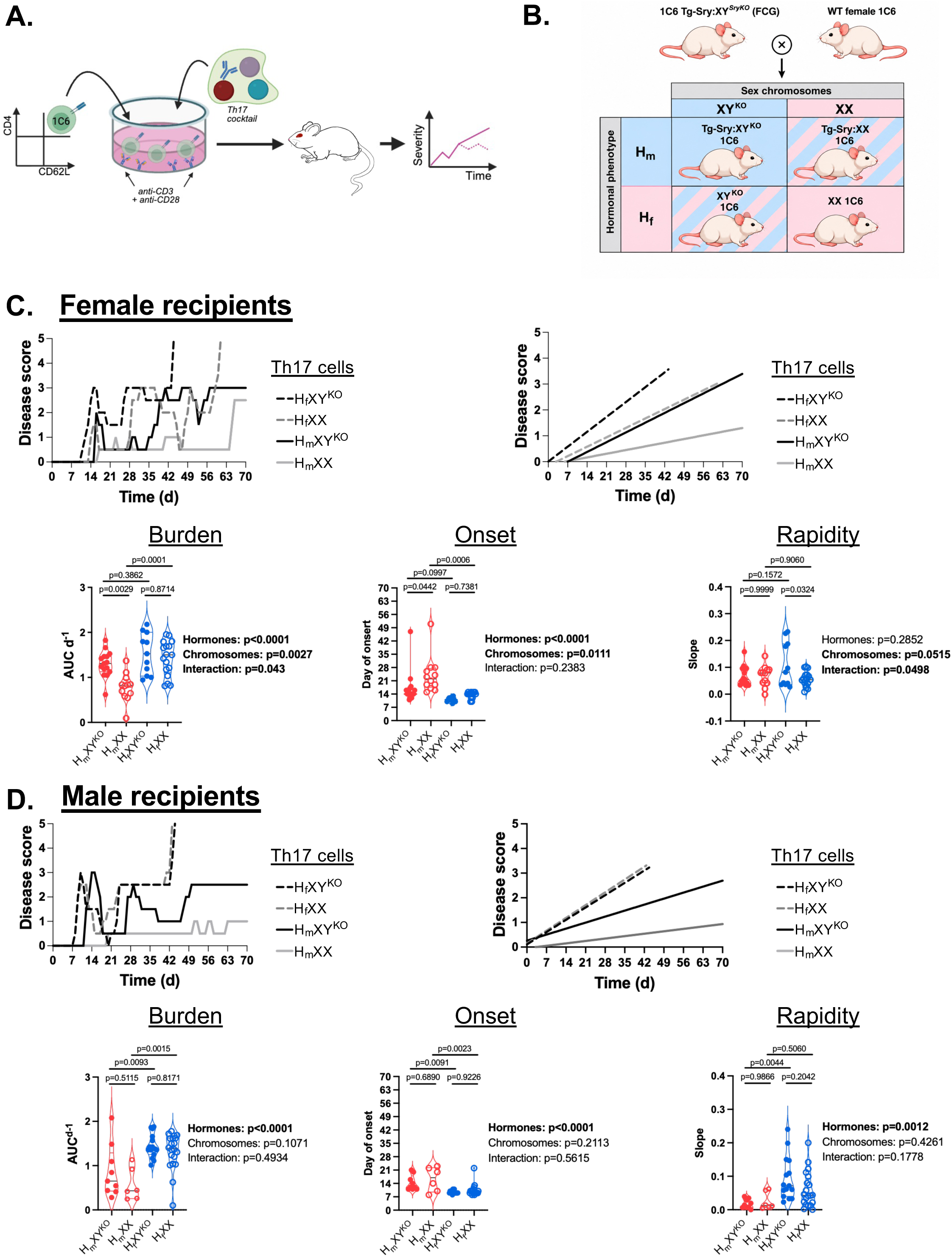
The effects of sex hormones and sex chromosomes on Th17 pathogenicity can be disaggregated. **A.** Schematic depicting 1C6 Th17 cell adoptive transfer model. NOD.*Scid* hosts receive 5×10^6^ Th17 cells. **B.** FCG:1C6 breeding strategy. **C.** Disease outcomes in female NOD.*Scid* recipients of FCG Th17 cells. *Top left*: representative disease curves for recipients of FCG Th17 cells. *Top right*: linear regression lines of the representative curves. *Bottom row*: quantification of disease burden (Area under curve per day; AUC d^-1^), disease onset and rapidity (slope of linear regression line). H_f_XY^KO^, n=11; H_f_XX, n=17; H_m_XY^KO^, n=15; H_m_XX, n=12. Summary data from 6 transfers. **D.** Disease outcomes in male NOD.*Scid* recipients of FCG Th17 cells. *Top left*: representative disease curves for recipients of FCG Th17 cells. *Top right*: linear regression lines of the representative curves. *Bottom row*: quantification of disease burden, onset and rapidity by hormonal and chromosomal variables. H_f_XY^KO^, n=15; H_f_XX, n=20; H_m_XY^KO^, n=9; H_m_XX, n=6. Summary data from 7 transfers. Hormone, chromosome and interaction effects were measured by two-way ANOVA. Pairwise comparisons were made by Tukey’s post-hoc test. Linked to Supplementary Figure 1.

To define the contributions of sex hormones and chromosomes to these phenotypes, we used the FCG model, which leverages the fact that the presence of *Sry* enforces male gonadal development while the absence of *Sry* results in a female developmental program. Double-transgenic FCG mice feature a knockout of endogenous *Sry* on the Y chromosome (Y*^SryKO^*) complemented by a knock-in of *Sry* on autosomal chromosome 3 (Tg-*Sry*). These mice are both chromosomally male and hormonally male, as the absence of endogenous *Sry* on Y is therefore complemented by ectopic *Sry*. When crossed to a WT female, two of the 4 possible genotypes are sex-concordant: male hormones (ectopic *Sry*, referred to herein as H_m_) with XY*^SryKO^* (referred to herein as XY^KO^), and female hormones (no ectopic *Sry,* or H_f_) and XX. By contrast, the other two are sex-discordant: male hormones (H_m_) with XX and female hormones (H_f_) with XY^KO^. We backcrossed the FCG line to the NOD genetic background and to the 1C6 strain. We subsequently bred FCG:1C6 mice to a non-FCG 1C6 female (schematic in Figure 1B); all progenies were therefore positive for the 1C6 transgene and were of one of the 4 possible sex genotypes.

Th17 cells from each genotype were first adoptively transferred to female NOD.*Scid* recipients. In agreement with our previous observations using FCG Th17 cells on a C57BL/6J (B6):NOD F1 genetic background^14^, we found that H_m_XX Th17 cells induced EAE with both a lower disease burden and delayed time of onset relative to gonadally-matched H_m_XY^KO^ Th17 cells. Disease burden and onset were also attenuated in recipients of H_m_XX Th17 cells relative to recipients of chromosomally-matched H_f_XX Th17 cells. Significant differences were not observed with respect to disease burden and onset when we compared recipients of H_m_XY^KO^ vs H_f_XY^KO^ Th17 cells, or recipients of H_f_XY^KO^ and H_f_XX Th17 cells (Figure 1C). A similar pattern emerged when we assessed the frequency of relapses: these were less frequent in recipients of H_m_XX Th17 cells versus mice that were transferred with either H_m_XY^KO^ or H_f_XX comparator cells (Supplementary Figure 1A). These data suggested that a male hormonal milieu, in combination with XX chromosomes, could potently suppress Th17-mediated CNS autoimmunity as measured by multiple parameters.

Curiously, in measuring the rapidity of disease accumulation in female recipient mice, we observed no differences between hosts of H_m_XX Th17 cells and the other groups. Rather, H_f_XY^KO^ Th17 cells induced more rapid disease than H_f_XX Th17 cells that were also gonadally female (Fig 1C), pointing to a pathogenic role for XY chromosomes as opposed to XX in a hormonally female context. To tease apart the relative contributions of sex hormones and sex chromosomes in regulating Th17 pathogenicity for each parameter, we went back to examine the main effects of hormones (H_m_ vs H_f_) and chromosomes (XX vs XY^KO^) separately. We found that sex hormones contributed significantly to the differences observed in disease burden and onset, with male hormones (H_m_) lessening pathogenicity. Sex chromosomes contributed significantly to differences in burden, onset and rapidity, with XY^KO^ augmenting disease (Figure 1C). Thus, in our model of Th17 cell-mediated pathogenicity, the effects of sex hormones and sex chromosomes could be uncoupled, and in fact could antagonize one another.

We next transferred FCG Th17 cells to male NOD.*Scid* recipients. Male sex hormones significantly attenuated Th17 burden and onset, with both H_m_XX and H_m_XY^KO^ Th17 cell recipients showing improvement on both disease parameters relative to their respective chromosomally-matched H_f_XX and H_f_XY^KO^ counterparts (Figure 1D). H_m_XY^KO^ recipients had reduced disease rapidity relative to H_f_XY^KO^ recipients, while no difference was detected in comparing mice receiving H_m_XX versus H_f_XX Th17 cells (Figure 1D). Conversely, H_m_XX Th17 cells elicited fewer relapses in host mice than H_f_XX counterparts, while no differences were observed between recipients of H_m_XY^KO^ and H_f_XY^KO^ Th17 cells (Supplementary Figure 1B). Thus, in male host mice, the effects of gonadal hormones on Th17 pathogenicity were dominant yet could be influenced by sex chromosomal complement. Together, our data from both female and male recipient animals showed that the effects of sex hormones and sex chromosomes on Th17 pathogenicity could be uncoupled: while male sex hormones improved disease parameters, XY chromosomes tended to worsen them.

### Sex hormones and chromosomes regulate distinct facets of the Th17 inflammatory phenotype in vivo

Having found that biological sex variables have profound influences on Th17 pathogenicity, we wanted to ascertain their influence on T cell phenotypes in vivo. Neither sex hormones nor sex chromosomes influenced the presence of transferred CD4^+^ T cells in the spleen (Supplementary Figure 2A) and brain (Supplementary Figure 2B) of recipient NOD.*Scid* mice. Thus, we next examined whether sex hormones and/or chromosomes impacted the production of inflammatory cytokines by transferred T cells. Gonadal sex potently impacted the pattern of cytokine production by brain-infiltrating Th17 cells. Female hormones were associated with reduced IL-17 and increased IFNγ and GM-CSF, a cytokine profile indicative of pathogenic Th17 cells in vivo^14,20–23^. Sex chromosomes were independently linked to differences in IFNγ, with XY Th17 cells producing more of this cytokine; our data thus supported a framework by which Th17 cell pathogenicity is shaped by both gonadal hormones and sex chromosomes.

### Sex hormones and sex chromosomes regulate Th17 proliferation and differentiation

To better understand the contribution of sex hormones and chromosomes to Th17 cell function, we next ascertained their impact on the initial steps of FCG Th17 cell differentiation in vitro. In first assessing parameters of cell proliferation, we found that H_f_XY^KO^ Th17 cells showed markedly greater expansion (Figure 2B) and per-cell division (Figure 2C) relative to hormone-matched H_f_XX or chromosome-matched H_m_XY^KO^ counterparts. Thus, the interaction of female gonadal hormones and XY chromosomes selectively enhanced the proliferative capacity of Th17 cells.

**Figure 2.**
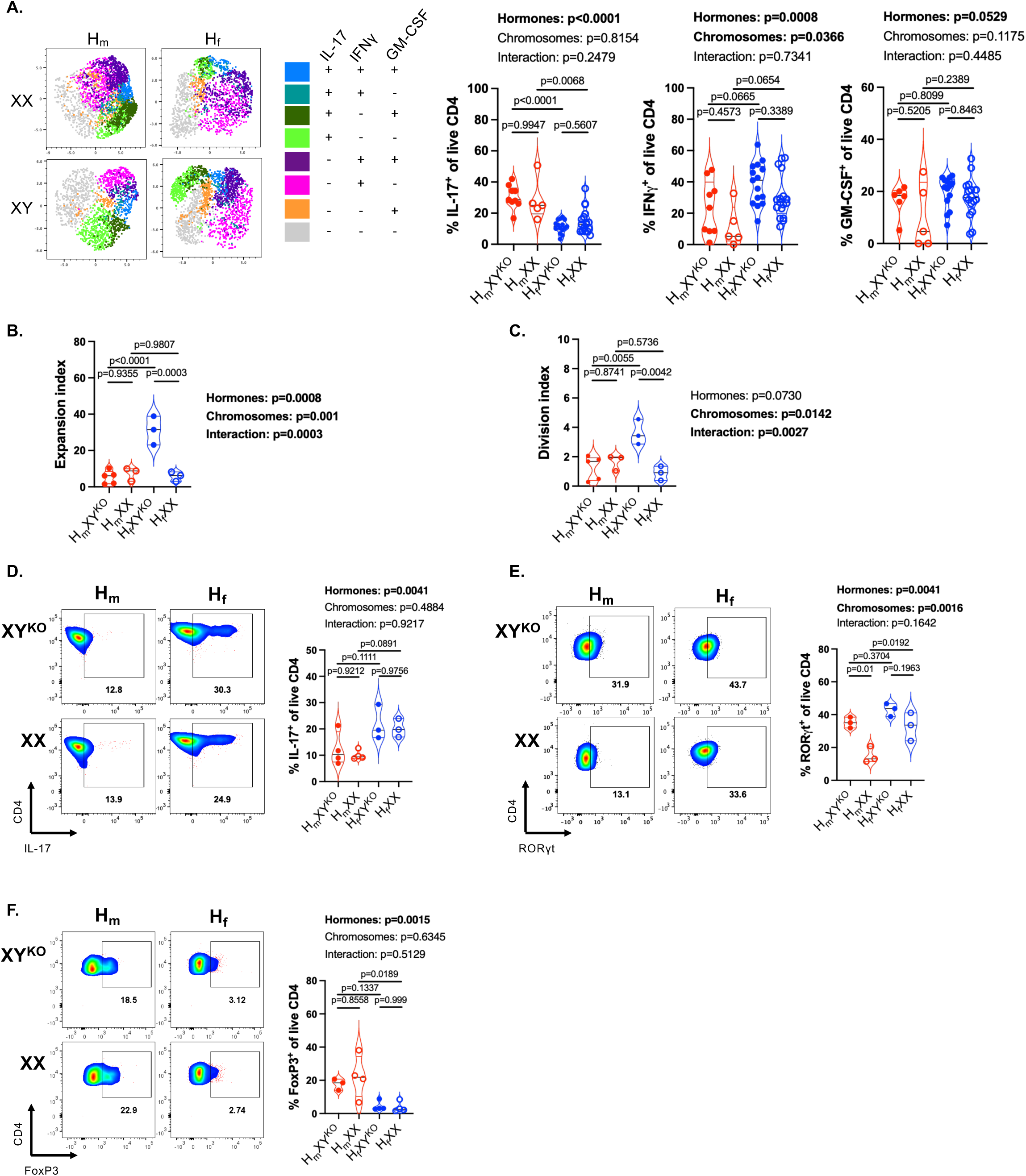
Regulation of Th17 function by hormonal and chromosomal sex. **A.** Representative UMAP plots and quantification of % IL-17, IFNγ and GM-CSF producers among brain-infiltrating FCG Th17 cells from male NOD.*Scid* recipients. H_f_XY^KO^, n=15; H_f_XX, n=17; H_m_XY^KO^, n=9; H_m_XX, n=5. **B, C.** Expansion index (**B**) and division index (**C**) were assessed from CellTrace Violet-labeled FCG Th17 cultures. H_m_XX n=3, H_m_XY^KO^ n=5, H_f_XX n=3, H_f_XY^KO^ n=3. **D.** Representative plots and quantification of IL-17A positivity from FCG Th17 cells. H_m_XX, n=3, H_m_XY^KO^, n=4, H_f_XX, n=3, H_f_XY^KO^, n=3. **E** Representative plots and quantification of RORγt positivity from FCG Th17 cells. H_m_XX, n=3, H_m_XY^KO^, n=3, H_f_XX, n=3, H_f_XY^KO^, n=3. **F.** Representative plots and quantification of FoxP3 positivity from FCG Th17 cells. H_m_XX, n=4, H_m_XY^KO^, n=3, H_f_XX, n=3, H_f_XY^KO^, n=3. Hormone, chromosome and interaction effects were measured by two-way ANOVA. Pairwise comparisons were made by Tukey’s post-hoc test. Each symbol is derived from an observation from a distinct mouse. Experiments in **B-F** were conducted 2-4 times. Linked to Supplementary Figure 2.

We then turned our attention to the impact of sex variables on markers of Th17 cell differentiation. Gonadal hormones but not sex chromosomes, contributed to differences in IL-17 with male hormones associated with reduced expression (Figure 2D). In examining RORγt expression, we found it to be markedly lower in H_m_XX Th17 cells than in either hormonally-matched H_m_XY^KO^ or chromosomally-matched H_f_XX Th17 cells (Figure 2E), suggesting that the combination of male sex hormones and XX chromosomes suppressed this master Th17 transcription factor ^24^. As it is well-established that Th17 cells and regulatory T cells (Treg) share a reciprocal relationship^25^, we subsequently examined whether sex variables could regulate FoxP3 in Th17 cells in vitro. Here, we found that male sex hormones contributed strongly to FoxP3 expression while sex chromosomes had no effect (Figure 2F). Together, these findings indicate that male sex hormones are consistently immunosuppressive across multiple aspects of the Th17 cell differentiation program, while the contribution of sex chromosomes is restricted to RORγt expression.

To ascertain whether sex hormones and/or sex chromosomes have an impact on T cell populations at baseline, we then examined thymocyte subpopulations in FCG:1C6 mice. Overall thymic cellularity was lowest in H_m_XX mice relative to H_m_XY^KO^ or H_f_XX counterparts (Supplementary Figure 2C). However, no sex-dependent differences were detected in the frequency of CD4^+^CD8^+^ double-positive (DP) thymocytes, nor were they observed for CD4^+^ single positive (SP) cells (Supplementary Figure 2D). Curiously, the combination of female gonadal hormones and XY chromosomes significantly augmented the frequency of CD8^+^ SP thymocytes, though the overall frequency of these cells remained <10% of all thymocytes (Supplementary Figure 2D). Turning to the spleen, we found that male sex hormones potently suppressed viable CD4^+^ T cell counts, while sex chromosomes had no effect (Supplementary Figure 2E).

### Androgen-related transcriptional programs and androgen receptor expression are linked to attenuated Th17 pathogenicity

We previously observed that adoptive transfer of WT male 1C6 Th17 cells causes widely divergent outcomes in recipient mice^14^, with approximately half developing severe EAE and the other half developing much milder EAE. As our data now revealed that gonadally male FCG Th17 cells display both reduced lineage commitment and pathogenicity, we asked whether differences in male sex hormone signaling might explain differential outcomes upon the transfer of male Th17 cells. We adoptively transferred WT male 1C6 Th17 cells to male NOD.*Scid* mice and monitored them daily for signs of EAE. For each mouse that attained ethical endpoints (M_sev_), we sacrificed a parallel recipient displaying mild disease (M_mild_). We then purified CD4^+^ T cells from brain and spleen from both sets of mice for transcriptional analysis. Using an established paradigm^26^, we defined brain-infiltrating T cells as pathogenic and splenic T cells as non-pathogenic (Figure 3A). In comparing the transcriptional profile of brain infiltrating male M_sev_ vs M_mild_ Th17 cells, we identified 913 differentially expressed genes (DEG) (29 upregulated in M_sev_, 884 downregulated; Supplementary Figure 3A). Interestingly, there were substantially more DEG (1775 total; 289 upregulated in M_sev_, 1486 downregulated) between splenic M_sev_ and M_mild_ Th17 cells (Supplementary Figure 3B), suggesting that differences in the pathogenic potential of male Th17 cells might be driven by transcriptional changes in the immune periphery. In line with this, we noted the complete absence of GO terms associated with DEG in the brain compartment. By contrast, 668 GO terms were associated with the comparison of M_sev_ vs M_mild_ splenic Th17 cells. These included “male gamete generation” (GO:0048232) and “spermatogenesis” (GO:0007283) in M_mild_ versus M_sev_ T cells (Fig 3B), both being processes that are critically dependent on androgens^27,28^. Furthermore, when we compared all brain-infiltrating Th17 cells (from M_sev_ and M_mild_ mice) with their splenic counterparts, we found that these two terms were enriched in cells taken from the spleen (Supplementary Figure 3C). Together, these analyses indicated that androgen-dependent mechanisms might limit the pathogenicity of male Th17 cells by promoting their sequestration outside of inflamed tissue.

**Figure 3.**
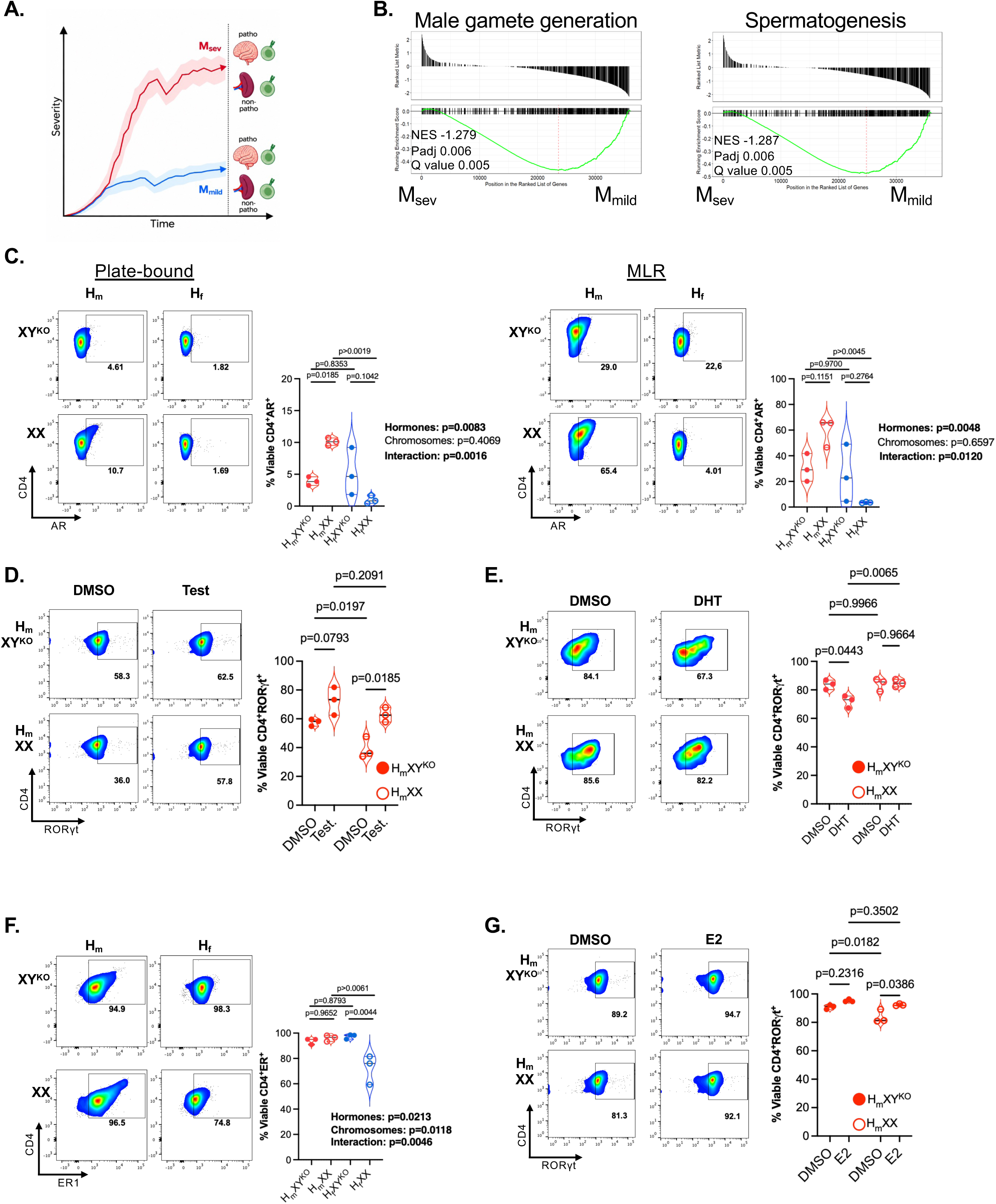
Androgen signaling reduces the pathogenicity of H_m_ Th17 cells. **A.** Schematic depicting the definitions of disease course severity (M_mild_ vs M_sev_) and T cell phenotype (non-pathogenic vs pathogenic). **B.** NES plots for the GO terms GO:0048232 (Male gamete generation) and GO:0007283 (Spermatogenesis), comparing the transcriptomes of splenic M_mild_ versus splenic M_severe_ Th17 cells. **C.** Representative plots and quantification of FCG:1C6 Th17 AR expression from plate-bound conditions (*left*, n= 3, each genotype) and MLRs (*right*, n= 3 each genotype). **D.** Representative plots and quantification of RORγt positivity from plate-bound stimulated H_m_XY^KO^ and H_m_XX 1C6 Th17 cells treated with 100 nM testosterone or DMSO (n= 3 each genotype). **E.** Representative plots and quantification of RORγt expression from plate-bound stimulated H_m_ H_m_XY^KO^ and H_m_XX 1C6 Th17 cells treated with 1 nM of DHT or DMSO (n= 3 each genotype). **F.** Representative plots and quantitation of ER1 expression from plate-bound stimulated FCG:1C6 Th17 cells (n=3 each genotype). **G.** Representative plots and quantification of RORγt positivity from plate-bound stimulated H_m_ 1C6 Th17 cells treated with 1 nM estradiol (E2) or DMSO (n= 3 each genotype). Each symbol represents a culture derived from an individual mouse. In **C** and **F**, hormone, chromosome and interaction effects were measured by two-way ANOVA. Pairwise comparisons were made by Tukey’s post-hoc test. In **D**, **E**, **G**, pairwise comparisons were made by Sidak’s multiple comparisons test. Experiments in **C-G** were conducted 2-4 times. Linked to Supplementary Figure 3.

Androgen receptor (AR) signaling suppresses the effector function of Th1 cells^29^ and restrains CD8^+^ T cell responses in cancer ^30,31^, and it was recently reported that testosterone can repress IL-17 production from Th17 cells^32^. We therefore turned to the FCG system to assess the potential contributions of sex hormones and chromosomes to the regulation of AR in Th17 cells. Under plate-bound stimulation conditions, in which T cells are stimulated with agonistic antibodies to CD3 and CD28 in the absence of other cell types, and mixed lymphocytic reaction (MLR) cultures, in which naïve FCG:1C6 CD4^+^ T cells were differentiated under Th17 conditions in the presence of both soluble anti-CD3 and irradiated splenocytes derived from the same mouse, we observed that male sex hormones were associated with increased AR positivity (Figure 3C). Interestingly, while we found no independent effect of sex chromosomes, we observed a significant interaction between sex hormones and sex chromosomes under both conditions, with greater AR positivity in H_m_XX relative to H_f_XX Th17 cells but no corresponding difference between H_m_XY^KO^ and H_f_XY^KO^ cells. Further, under plate-bound conditions alone, AR positivity was higher in H_m_XX Th17 cells as compared to H_m_XY^KO^ cells. Thus, the combination of male sex hormones and XX chromosomes was associated with both increased AR positivity from Th17 cells as well as with attenuated pathogenicity of these cells.

### Androgenic and estrogenic regulation of Th17 differentiation is influenced by sex chromosome complement

We next assessed the impact of AR signaling on H_m_ FCG Th17 cell differentiation. Given the predicted inhibitory effect of androgen signaling on Th17 function, we were surprised to observe that testosterone increased expression of RORγt, most notably from H_m_XX Th17 cells (Figure 3D). Interestingly, however, we found that dihydrotestosterone (DHT), a testosterone metabolite and potent androgen, markedly reduced RORγt expression from H_m_XY^KO^ Th17 cells yet had no effect on H_m_XX (Figure 3E).

As testosterone^33^, but not DHT^34^, can be aromatized to generate estrogen, we asked whether the dichotomous effects of these two androgens on RORγt expression might be explained by a countervailing stimulatory effect of estrogen. Indeed, we found that the estrogen receptor (ER1) is highly expressed on both hormonally male and hormonally female FCG Th17 cells (Figure 3F). Furthermore, and in line with what we had observed upon treatment with testosterone, estradiol increased the expression of RORγt from H_m_XX, but not H_m_XY^KO^, Th17 cells (Figure 3G). Together, these findings indicate that gonadal hormones regulate Th17 cell differentiation, but that their effects are strongly influenced by sex chromosomes. While DHT suppressed RORγt expression selectively in H_m_XY^KO^ cells, estradiol increased its expression specifically in H_m_XX cells, suggesting that XX and XY chromosomes confer differential responsiveness to androgenic and estrogenic signals.

### Androgen mediated Wnt signaling pathway drives reduced pathogenicity of male Th17 cells

We next wanted to evaluate biological processes that might associate with the AR signaling pathway to modulate the pathogenicity of male Th17 cells. We noted that the GO term *“Wnt signaling pathway”* (GO:0016055) was enriched in the transcriptome of male splenic Th17 cells as compared to male brain-infiltrating cells (Supplementary Figure 4A), and in splenic M_mild_ Th17 cells relative to splenic M_sev_ Th17 cells (Figure 4A). Turning once again to the FCG system, we then observed that male gonadal hormones were associated with increased expression of the Wnt signal transducer protein β-catenin (Figure 4B). The Wnt effector molecule TCF-1 was also increased in H_m_XY^KO^ Th17 cells relative to H_f_XY comparators, although no differences were observed between Th17 cells derived from H_m_XX and H_f_XX mice (Figure 4C).

**Figure 4.**
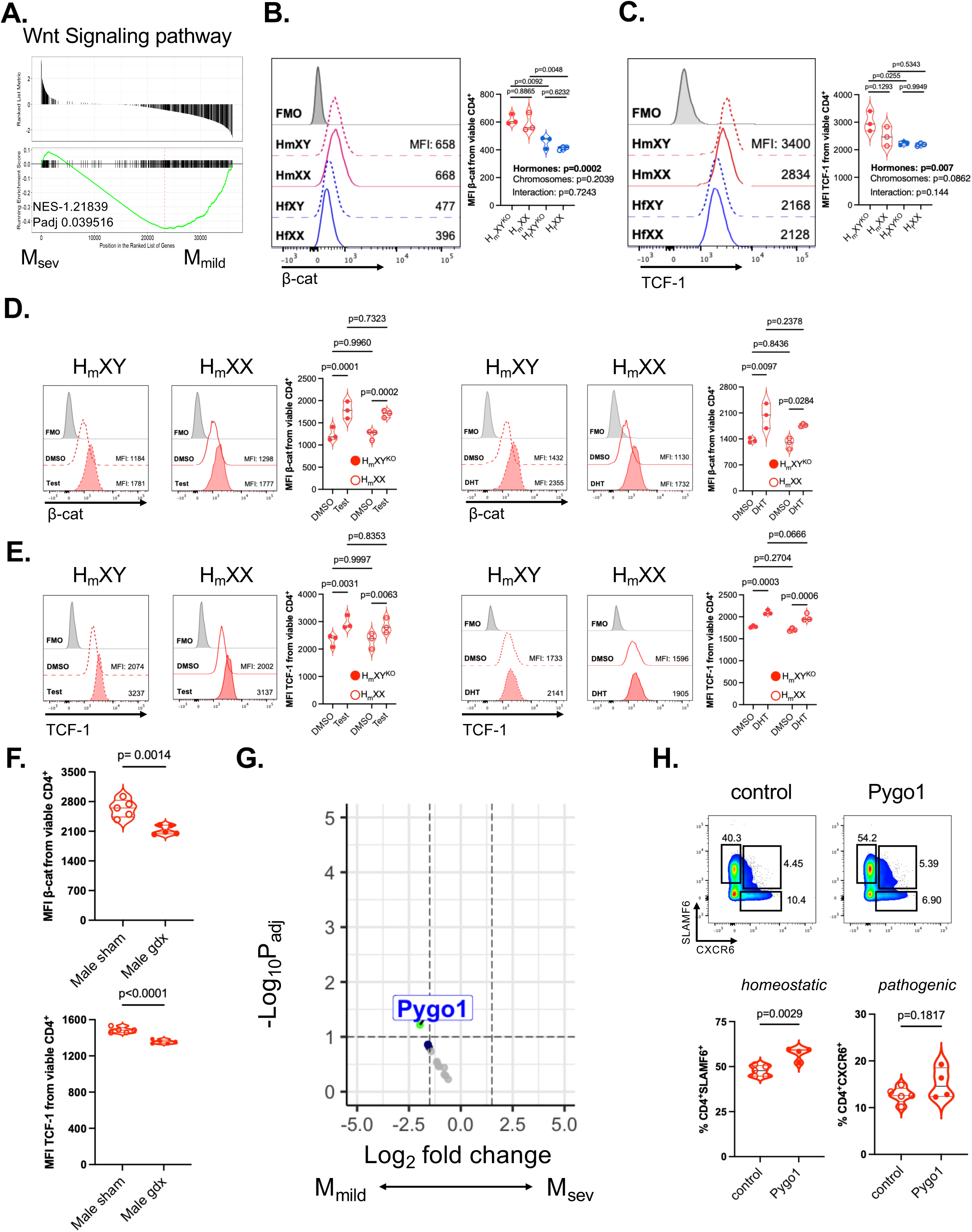
Androgen-associated Wnt signaling promotes a homeostatic Th17 phenotype. **A.** GSE plots for GO:0016055 (Wnt signaling pathway) comparing M_severe_ versus M_mild_ splenic Th17 cells. **B.** Representative histograms and quantification for β-catenin expression from MLR-stimulated FCG Th17 cells (n=3 of each genotype). **C.** Representative histograms and quantification for TCF-1 expression from MLR-stimulated FCG Th17 cells (n=3 of each genotype). **D.** Representative histograms and quantification of β-catenin expression from H_m_ Th17 cells (n=3 each genotype) treated with testosterone (100nM) or DHT (1nM). **E.** Representative histograms and quantification of TCF-1 expression from H_m_ Th17 cells (n=3 each genotype) treated with testosterone (100nM) or DHT (1nM). **F.** Quantification of β-catenin and TCF-1 expression from sham-operated or gonadectomized WT male 1C6 Th17 cells stimulated under plate-bound conditions (n=5 each sex). **G.** Volcano plot of expression differences of genes tagged with GO terms “spermatogenesis”, “male gamete generation and “Wnt signaling pathway”, between M_mild_ and M_sev_ splenic Th17 cells. **H.** WT male 1C6 Th17 cells were transduced with control- or Pygo1-expressing retrovirus and were adoptively transferred to male NOD.*Scid* recipients (n=5, control; n=4, Pygo-1). Expression of SLAMF6 and CXCR6 was assessed 10 days after transfer from splenic CD4^+^GFP^+^ cells. Each symbol represents a culture derived from an individual mouse. **A, G**, results were analyzed by the Benjamini-Hochberg procedure; **B, C**, hormone, chromosome and interaction effects were measured by two-way ANOVA and pairwise comparisons were made by Tukey’s post-hoc test; **D, E**, Sidak’s multiple comparisons test after repeated measures ANOVA; **F, H**, unpaired *t*-test. P_adj_ and log_2_ fold change cut-off in **G** were set at 0.1 and 1.5, respectively. Experiments in **B-E** were conducted 2-3 times. Linked to Supplementary Figure 4.

TCF-1 drives a homeostatic and non-pathogenic phenotype in Th17 cells^35^, and represses the severity of Th17 cell-mediated EAE^35,36^. Our findings thus raised the possibility that AR signaling might diminish Th17 pathogenicity by enhancing the Wnt pathway. To test this, we treated FCG H_m_ Th17 cells with testosterone or DHT, finding that both AR agonists increased the expression of β-catenin (Figure 4D) as well as of TCF-1 (Figure 4E) from both H_m_XY^KO^ and H_m_XX Th17 cells. Next, to extend our findings to the WT male setting, we generated Th17 cells from the spleens of gonadectomized or sham-operated male 1C6 mice and studied the expression of both β-catenin and TCF-1. Both proteins were downregulated in cultures from gonadectomized male mice (Figure 4F), again demonstrating that androgens promote Wnt activation in Th17 cells.

We next wanted to identify genes linked to both male sex hormone signaling and Wnt signaling that might diminish Th17 pathogenicity. As the GO terms “spermatogenesis”, “male gamete generation” (Figure 3B) and “Wnt signaling pathway” were all enriched in M_mild__splenic Th17 cells (Figure 4A), we focused on the 23 genes that were tagged with all 3 terms, reasoning that this might reveal novel molecules relevant to Th17 function that were co-regulated by both male sex hormones and Wnt (Supplementary Figure 4B). Of these 23 genes, *Pygo1* was the only one to be differentially regulated between M_mild_ and M_sev_ splenic Th17 cells, with greater expression in M_mild_ (Figure 4G). Pygo1 is a transcriptional co-activator of the Wnt signaling pathway^37^, leading us to consider whether it might promote a homeostatic Th17 phenotype. Indeed, while associated with kidney development^37^, cardiac hypertrophy^38^ and cancer cell invasiveness^39^, a role for Pygo1 in T cell function has not previously been described. We therefore retrovirally overexpressed Pygo1 (Pygo1-OE) in male and female WT 1C6 CD4+ T cells while differentiating them under Th17 conditions. While we expected that ectopic Pygo1 would restrain Th17 cell differentiation, we observed, surprisingly, that it significantly increased RORγt expression from male Th17 cells relative to control transduced cells (Supplementary Figure 4C). However, recalling that TCF-1^+^ Th17 cells display enhanced persistence^40^, we next asked whether Pygo1 might specifically favor the development of this homeostatic Th17 cell subset. Indeed, Pygo1-OE increased expression of TCF-1 in male RORγt^+^, but not RORγt^-^ CD4^+^ T cells (Supplementary Figure 4D). This effect was not observed in Pygo1-OE female Th17 cells (Supplementary Figure 4E), suggesting that Pygo1 expands TCF-1^+^ Th17 cells in a sex-dependent manner.

In the in vivo setting, homeostatic Th17 cells are marked by SLAMF6, whose expression is in turn promoted by TCF-1. As ectopic Pygo1 augments TCF1 expression in male Th17 cells, we asked whether it may also favor the generation of a homeostatic phenotype in vivo. We therefore adoptively transferred Pygo1-OE or control-transduced male 1C6 Th17 cells to male NOD.*Scid* hosts. After 10 days, we isolated splenic T cells from recipient animals and found that Pygo1-OE Th17 cells displayed a robust increase in SLAMF6 expression relative to control cells. By contrast, we observed no differences in the frequency of Th17 cells expressing the inflammatory marker CXCR6 (Figure 4H). Taken together, our data showed that male gonadal hormones positively regulate expression of the core Wnt pathway signaling components β-cat and TCF-1. Further, the Wnt transcriptional co-activator Pygo1 is upregulated in non-pathogenic male Th17 cells and promotes a homeostatic phenotype.

### X-linked SUV39H1 negatively regulates Th17 responses

We had shown that XY chromosomal complement could augment Th17 cell pathogenicity (Figure 1). A recent study had reported that the mutant Y*^SryKO^* chromosome of FCG mice carries a 3.2 Mb X-chromosome translocation containing 9 genes^41^, one of which is the well-established pattern recognition receptor and inflammatory mediator *Tlr7* ^42^; however, we established that TLR7 is only expressed at low levels on FCG Th17 cells and is not regulated by either sex hormones or sex chromosomes (Figure 5A). To assess whether the other genes in the translocated region might be upregulated in cells carrying the transgenic Y*^SryKO^* chromosome, we conducted RNAseq analysis of FCG Th17 cells and compared the expression of X-encoded genes between chromosomally female (H_f_XX, H_m_XX) and chromosomally male (H_f_XY, H_m_XY) samples. Of the nine genes in the translocated region, four (*Prps2*, *Tmsb4x*, *Msl3* and *Hccs*) showed significantly increased expression in XY cells (Supplementary Table 1). We examined the protein expression of PRPS2 and TMSB4X, the two genes displaying the greatest fold changes and found that neither showed differential expression between H_m_XX and H_m_XY, or H_f_XX and H_f_XY, Th17 cells (Supplementary Figure 5A). Thus, our findings suggest that the increased pathogenicity of XY Th17 cells is unlikely to be driven by increased protein expression of the most transcriptionally dysregulated genes within the reported translocated region.

**Figure 5.**
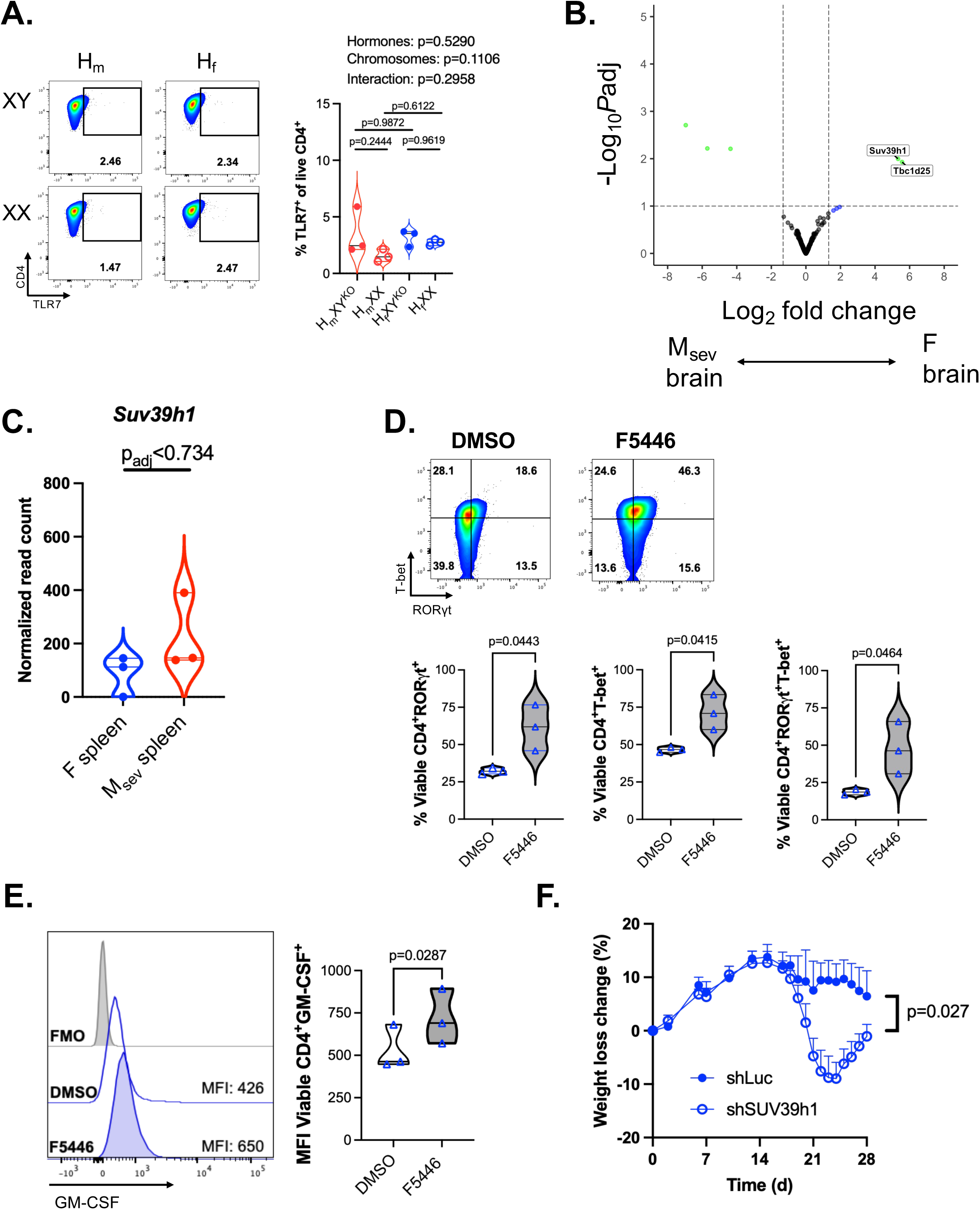
X-linked *Suv39H1* suppresses female Th17 cell responses. **A.** Representative plots and quantification for TLR7 expression from plate-bound-stimulated FCG Th17 cells (n=3 of each genotype). **B.** Volcano plot depicting differential expression of X-linked genes between brain-infiltrating female (F) and M_sev_ 1C6 Th17 cells (n=3 each). **C.** *Suv39h1* expression in splenic F versus M_sev_ Th17 cells. **D.** Female 1C6 Th17 cells were treated with 50 nM F5446 (n=3) or DMSO (n=3) for 5 days and expression of RORγt and T-bet was assessed on CD4^+^ T cells by flow cytometry. **E.** 1C6 female Th17 cells were treated with 50 nM F5446 (n=3) or DMSO (n=3) for 5 days and expression of GM-CSF was assessed on CD4^+^ T cells by flow cytometry. **F.** Female 1C6 Th17 cells were transduced with SUV39h1-shRNA- or luciferase-shRNA-encoding retrovirus and were adoptively transferred to female NOD.*Scid* recipients (2 × 10^6^ per mouse). Data presented as percentage weight loss relative to weight at the time of adoptive transfer. n=5 SUV39h1-shRNA, n=4 luciferase (Luc)-shRNA. Each symbol represents a culture or analysis derived from an individual mouse. In **A**, Hormone, chromosome and interaction effects were measured by two-way ANOVA and pairwise comparisons were made by Tukey’s post-hoc test. In **B**, **C**, the Benjamini-Hochberg procedure was used; P_adj_ and log_2_ fold change cut-off in **B** were set at 0.1 and 1.5, respectively. Ratio paired *t*-test (**D, E**), two-way ANOVA (**F**). Experiments in D, E were conduced two times. Linked to Supplementary Figure 5.

In therian mammals, most genes on the X chromosome are expressed from a single copy in female cells, with one X chromosome being silenced via effects mediated by the long-coding RNA *Xist* ^43^. However, second-allele silencing is far from a complete process, and a substantial frequency of X-chromosome genes can be expressed from both copies in female cells in a tissue- and temporally-specific manner^11^. Indeed, X-encoded molecules such as Kdm6a^44^ and Kdm5c^14^ can influence disease outcomes in T cell-driven CNS autoimmunity. We therefore sought to determine, in an unbiased manner, whether variation in the expression of X-linked genes could account for the differences in disease severity observed between female and male recipients of 1C6 Th17 cells. We first compared the expression of X-linked genes between the transcriptome of adoptively transferred, brain-infiltrating, female 1C6 Th17 cells and that of brain-infiltrating M_sev_ cells. Two X-encoded genes, *Suv39h1* and *Tbc1d25*, displayed increased expression in female cells (Figure 5B). SUV39H1 is a histone methyltransferase previously implicated in driving Th2 lineage commitment at the expense of the Th1 cell fate^45^. We observed that *Suv39h1* was similarly expressed in female and M_sev_ Th17 cells isolated from the spleens of recipient mice (Figure 5C), indicating that it was differentially regulated specifically in brain-infiltrating cells. We thus speculated that SUV39H1 might regulate the inflammatory functions of Th17 cells.

CNS-infiltrating Th17 cells express the Th1 transcription factor T-bet, which is essential to their pathogenicity ^46,47^. As SUV39H1 represses T-bet expression in Th2 cells^45^, we asked whether it might have a similar function in female Th17 cells. We treated WT female 1C6 Th17 cells with F5446, a SUV39h1-specific inhibitor^48^, or with vehicle. Strikingly, F5446 treatment increased the expression of both RORγt and T-bet, as well as the frequency of RORγt^+^T-bet^+^ double-positive cells (Figure 5D). Further, expression of pathogenic GM-CSF was increased in female Th17 cells upon F5446 treatment (Figure 5E).

To test whether SUV39H1 represses the pathogenicity of female 1C6 Th17 cells, we transduced WT female 1C6 Th17 cells with either sh*Suv39h1*- or control (luciferase)-shRNA-expressing retrovirus that we subsequently transferred to female NOD.*Scid* recipients as we performed previously^49^. SUV39H1 depletion in Th17 cells elicited significant weight loss in recipient mice (Fig. 5F), a well-established correlate of pathology in EAE^50,51^ and in T cell adoptive transfer models^52,53^. Our data thus showed that sex chromosome-driven differences in Th17 cell-induced disease were not driven by genes present in the previously described X-Y translocation on the genetically modified Y*^SryKO^* chromosome. Further, they revealed the histone methyltransferase SUV39h1 as a regulator of inflammatory Th17 cell function in vitro and in vivo.

### Hormonally and chromosomally female H_f_XX Th17 cells are highly pathogenic due to reduced expression of Runx2

We had previously shown that female 1C6 Th17 cells transfer mild disease^14^. We were therefore struck by the observation that H_f_XX 1C6 Th17 cells induced disease of relatively strong severity (Figure 1C, D), even though H_f_XX mice are both hormonally and chromosomally female. Indeed, a side-by-side comparison of EAE induced by WT female 1C6 versus H_f_XX:1C6 Th17 cells revealed that the latter elicited a strikingly higher disease burden (Figure 6A). Next, we gonadectomized H_f_XX:1C6 mice and generated Th17 cells that we adoptively transferred in parallel with recipients of Th17 cells from sham-operated H_f_XX:1C6 Th17 donors. We observed no differences between the two groups (Supplementary Figure 6A), suggesting that the unexpected pathogenicity of H_f_XX:1C6 Th17 cells was not due to alterations in circulating sex hormones in donor mice. Additionally, we observed no differences in expression of *Xist* expression between H_f_XX:1C6 and female 1C6 Th17 cells, suggesting that differences in the pathogenicity of cells from these two genotypes was not due to differences in X chromosome inactivation (Supplementary Figure 6B).

**Figure 6.**
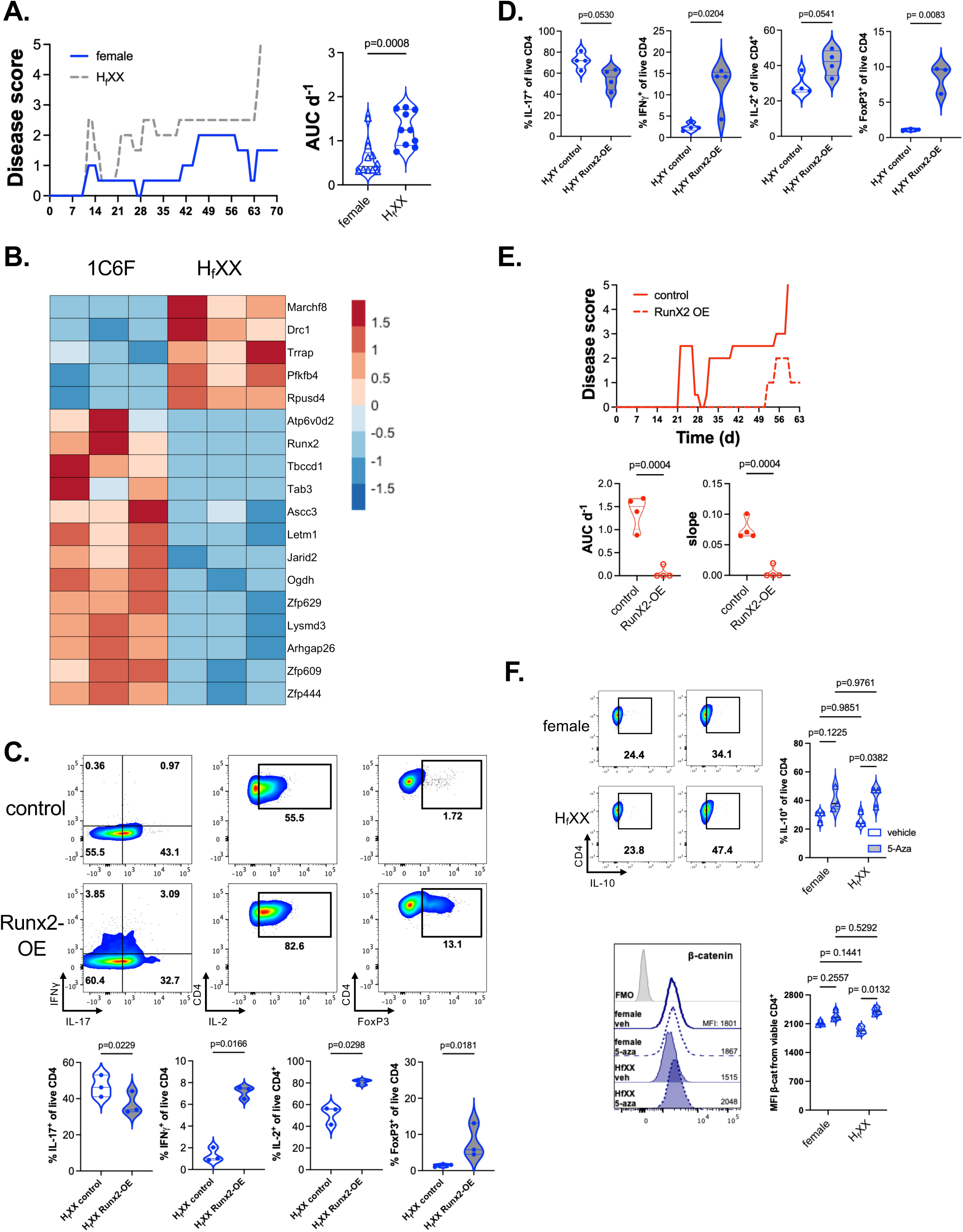
H_f_XX Th17 cells are more pathogenic than female Th17 cells. **A.** WT female 1C6 or H_f_XX 1C6 Th17 cells were adoptively transferred to male NOD.Scid mice that were monitored for signs of EAE. *Left*, representative disease curves and linear regression lines. *Right*, quantification of disease burden. n=10, each group of recipients. Pooled from two adoptive transfer experiments. **B.** Heatmap depicting DEGs between H_f_XX (n=3) and 1C6 female (n=3) Th17 cells. Padj and log_2_ fold-change were set at 0.05 and 0.4, respectively. **C.** H_f_XX 1C6 Th17 cells were transduced with control- or Runx2-expressing retrovirus. Representative plots and quantification of expression of the indicated cytokines or of FoxP3 after 5 days of culture (n=3 each condition). **D.** H_f_XY 1C6 Th17 cells were transduced with control- or Runx2-expressing retrovirus. Representative plots and quantification of expression of the indicated cytokines (n=4 each condition) or of FoxP3 (n=3 each condition) after 5 days of culture. **E.** H_f_XY 1C6 Th17 cells were transduced with control- or Runx2-expressing retrovirus and were adoptively transferred (1 × 10^6^ per mouse) to male NOD.*Scid* recipients (n=4 each group). *Top*, representative disease curves. *Bottom*, quantification of disease burden and rapidity. **F.** WT female 1C6 and H_f_XX 1C6 Th17 cultures (n=3 each) were treated with 0.5 μM 5-azacytidine or DMSO vehicle. Representative plots and quantification of expression of IL-10 and β-catenin after 5 days of culture. Each symbol represents a culture derived from an individual mouse. Results were analyzed by unpaired t-test (**A, E**), ratio paired t-test (**C, D**) or Sidak’s multiple comparisons test (**F**). Experiments in C, D, F were conducted two times. Linked to Supplementary Figure 6.

Reasoning that differences in gene expression would ultimately underpin this difference in pathogenicity, we conducted bulk RNA-seq on H_f_XX:1C6 and WT female 1C6 Th17 cells after 5 days of culture. We identified 18 DEG (Figure 6B), a number of which have putative roles in regulating gene expression: the histone modifier *Trrap* ^54^, the transcription factors *Runx2* ^55^ and *Zfp609* ^56^ or DNA binders such as *Assc3* ^57^ and *Jarid2* ^58^. *Runx2* attracted our specific interest as it plays diverse roles in dictating T lymphocyte cell fate: it regulates the transition of thymocytes from the DN3 to DN4 cell fates ^59^, promotes the long-term in vivo persistence of memory CD8^+^ T cells ^60^ and is a transcriptional repressor of T follicular helper (Tfh) cells^61^. As Runx2 was downregulated in pathogenic H_f_XX:1C6 Th17 cells, we therefore asked whether it might repress Th17 function. We generated a Runx2-OE retroviral construct and used it to transduce H_f_XX Th17 cells in vitro. Runx2-OE reduced the expression of IL-17 from H_f_XX Th17 cells while augmenting the expression of IFNγ as well as of IL-2, a well-described antagonist of Th17 differentiation^62,63^; moreover, Runx2-OE increased expression of FoxP3 (Figure 6C). To examine whether these effects were generalizable, or unique to the H_f_XX context, we expressed Runx2-OE in H_f_XY Th17 cells, which also show robust Th17 responses in vitro (Figure 2E). We found that Runx2-OE H_f_XY Th17 cells again increased IFNγ and FoxP3 expression relative to control comparators (Figure 6D). These data indicated that Runx2 diminishes Th17 lineage fidelity and might promote a regulatory phenotype in Th17 cells.

Though the increase in FoxP3 expression in Runx2-OE-transduced cells suggested that Runx2 reduces Th17 pathogenicity, their elevated production of IFNγ raised the alternate possibility that Runx2 might worsen Th17-mediated disease. To distinguish between these possibilities, we transduced H_f_XY^KO^ Th17 cells, which elicit severe EAE, with Runx2-OE or control RV. We found that ectopic expression of Runx2 nearly completely abrogated EAE upon adoptive transfer of these cells (Figure 6E). Together, our data showed that Runx2 is among a small cluster of genes whose expression is reduced in H_f_XX versus WT female Th17 cells, and that Runx2 powerfully suppresses Th17 pathogenicity. Thus, the increased pathogenicity of H_f_XX Th17 cells relative to WT female Th17 cells may be due in part to reduced Runx2 expression.

### DNA methylation underpins a pro-inflammatory phenotype in H_f_XX Th17 cells

H_f_XX mice bear neither FCG modified allele and differ from WT females only in that they have an H_m_XY^KO^ sire (Figure 1B). Notably, we observed no differences in sperm concentration, live sperm percentage and motility between H_m_XY^KO^ and WT males (Supplementary Figure 6C). Further, we found no differences in circulating testosterone and progesterone between H_m_XY^KO^ and WT males (Supplementary Figure 6D), while levels of E1, E2, E3 and DHT were undetectable.

DNA methylation is the best-described mechanism of epigenetic parental inheritance^64^ and is linked to differences in gene expression in T cells ^65^. We therefore asked whether this might explain the enhanced inflammatory profile of H_f_XX Th17 cells by treating them with the 5’-aza-2’-deoxycytidine (5-Aza), a DNA methyltransferase inhibitor that can derepress the expression of methylated genes. 5-Aza had no impact on the presence of IL-17^+^ cells (Supplementary Figure 6E); however, it increased the frequency of H_f_XX cells, but not WT female cells, that produced the immunosuppressive cytokine IL-10 (Figure 6F). The percentage of RORγt^+^ Th17 cells was unaltered by 5-Aza treatment; however, in line with previous work^66^, 5-Aza augmented FoxP3^+^ cells among WT as well as H_f_XX Th17 cell cultures (Supp Fig 6E). Intriguingly, 5-Aza increased the expression of β-catenin in H_f_XX, but not WT female, Th17 cells (Fig 6F). Altogether, these data indicate that DNA methylation represses the expression of multiple immune inhibitory genes in H_f_XX Th17 cells.

## Discussion

While MS incidence is skewed towards females, males with MS more rapidly attain signposts of advanced disability, such as the need to use a walking implement, and are more prone to cognitive decline ^12^. Curiously, a similar pattern has been noted in lupus^67^ and systemic sclerosis^68^, whereby disease incidence is higher in females while severity is increased in males. A deeper understanding of sex-specific differences in T cell function would shed light on these conflicting dichotomies. Th17 cells are of particular interest, as multiple Th17-dependent secreted inflammatory mediators are increased in the plasma of people with SPMS versus those with RRMS^69^. Further, in an experimental model, Th17 cells are essential to the subpial cortical injury that characterizes progressive MS^70^. Importantly, several human studies have reported that Th17 responses are augmented in males^71–73^, while in experimental models of myocarditis^10,74^ and arthritis^75^, male sex favors Th17-mediated disease. As we had previously shown that male 1C6 Th17 cells had the potential to elicit severe progressive CNS autoimmunity in a substantial proportion of recipient mice, we therefore wanted to tease apart the relative contributions of sex hormones and chromosomes to this phenotype. In the current study, we report that male sex hormones lessen the severity of Th17-mediated progressive EAE while XY chromosomes are associated with worsened disease.

We find that androgen-associated gene signatures are enriched in male 1C6 Th17 cells that invoke mild EAE as opposed to male Th17 cells that elicit severe disease. It has been reported that low testosterone levels may correlate with worsened outcomes in men with MS^76^. Further, a pilot clinical trial showed that exogenous testosterone could improve cognitive and imaging outcomes in men with MS^77,78^. The salutary impact of testosterone in MS has been attributed to its neuroprotective effects^79^. However, androgens have well-established immune suppressive effects on CD8^+^ T cells^30,31,80^ and they suppress the proliferation of CD4^+^ T cells ^81^ while promoting IL-10 production ^82^. Further, an elegant recent study demonstrated that testosterone suppress IL-17 production from human and mouse Th17 cells by inhibiting the binding of RORγt to the *Il17* and *Il23r* promoters^32^. In this study, we show that male sex hormones increase AR positivity in Th17 cells, an effect that is particularly pronounced in cells bearing XX chromosomes. Furthermore, DHT suppressed RORγt expression in H_m_XY^KO^, but not H_m_XX, Th17 cells, revealing a chromosome-dependent response to androgen signaling. Interestingly, ER1 was highly expressed on hormonally male Th17 cells, and estradiol increased RORγt expression specifically in H_m_XX cells. These findings suggest that aromatization to estrogen may explain the paradoxical increase in RORγt following testosterone treatment, while also revealing a critical influence of sex chromosome complement on estrogen responsiveness. Estrogen receptors are present on T cells from men^83,84^ and their expression is highly sensitive to environmental conditions^83^. It is possible that differential estrogen receptor expression on male Th17 cells may influence their effector function and thereby contribute to sex-specific differences in multiple sclerosis progression; this will be an important area for future study.

We demonstrate that mildly pathogenic male Th17 cells upregulate Wnt signaling-associated genes and that exogenous androgen increases expression of β-catenin and TCF1. These observations are notable because it was recently shown that TCF1 drives a homeostatic, rather than pathogenic, Th17 phenotype^35^. β-catenin physically interacts with AR^85–88^ and augments expression of AR-dependent genes in prostate cancer cells^88,89^. However, AR binding can increase β-catenin translocation to the nucleus ^85,88^, suggesting the opposite is also true and that androgen signaling can increase Wnt activity. Together, our data link male sex hormones to augmented Wnt signaling and thereby to diminished pathogenicity of Th17 cells.

Using B6:NOD F1 mice, we previously showed that H_m_XY 1C6 Th17 cells caused EAE of greater severity than H_m_XX counterparts^14^. Here, using a line fully backcrossed to NOD, we extend those findings by demonstrating that XY chromosomes can augment Th17 pathogenicity, although their effects depend on the hormonal context and sex of the recipient. Previous studies have indicated that the contribution of sex chromosomes to autoimmunity are nuanced and dependent on both genetic background and cellular compartment. SJL/J-strain FCG XY mice are less susceptible to peptide-immunization EAE, and to pristane-induced lupus, relative to XX comparators. By contrast, no differences between XX and XY mice are observed in MOG_[35-55]_ B6 EAE^90^. Of note, SJL/J XY T cells showed more potent proliferation and cytokine responses to antigenic recall, providing early evidence that male chromosomal complement might promote inflammation in a T cell-intrinsic manner^7^. It should be noted that this latter work was conducted prior to the characterization of Th17 cells as a distinct T cell effector lineage and thus IL-17 production was not assessed. Overall, these findings and our own highlight the importance of using diverse models to tease apart the complex contributions of sex chromosomes to immune responses.

As many as 20% of X-linked genes can be expressed in two copies in females in a tissue-specific manner^6^. Having identified a pathogenic contribution of XY chromosomes, we explored the possibility that X-linked immunomodulatory genes might restrain inflammatory responses in Th17 cells. Transcriptional analysis of male 1C6 Th17 cells that cause severe disease, versus female counterparts, revealed Suv39h1 as an X-linked gene upregulated in the latter. Pharmacological inhibition of Suv39h1 augmented the inflammatory phenotype of female 1C6 Th17 cells both in vitro and in vivo. Suv39h1 is a critical enforcer of the Th2 phenotype^45^ and can repress *Il2* transcription in anergic T cells^91^, indicating its importance in dictating effector T cell lineage. More broadly, these findings contribute to the accumulating evidence that variable expression or activity of X-linked genes can impact T cell function. The incidence of progressive MS increases after andropause^92^ and menopause^93^, suggesting that sex hormones might mask an underlying propensity for severe, chronic neuroinflammatory responses that is in part dictated by factors encoded on sex chromosomes. For example, we previously showed that the X-linked H3K4 demethylase Kdm5c/Jarid1c represses the pathogenicity of female 1C6 Th17 cells and is downregulated in T cells from people with MS^14^. By contrast, the related gene Kdm6a, which is also located on the X chromosome, promotes T cell-dependent neuroinflammation. The relative balance of X-linked factors such as Kdm5c and Kdm6a^94^, as well as Suv39h1, could thus dictate pathogenic outcomes driven by T cells when sex hormones are limited or absent.

The most frequently used FCG strain (Jackson Laboratories #010905) was recently described as carrying a translocation of a 3.2 Mb segment of X chromosomal DNA, containing 9 genes, to the transgenically modified Y chromosome^41^. We thus compared the transcript and protein expression of these genes between hormonally matched XX and XY Th17 cells. While several genes within the translocated region showed increased transcript expression in XY cells, we did not observe corresponding differences in protein expression for those exhibiting the greatest fold changes. These findings suggest that increased expression of these two genes is unlikely to account for the chromosomal sex differences observed in our model, although contributions from other genes within the translocated region cannot be excluded. The reported X-Y translocation represents an unexpected variable that will need to be accounted for carefully in the interpretation of data generated with this FCG strain. Here, we provide a possible roadmap for addressing this important consideration going forward.

The unexpectedly higher pathogenicity of H_f_XX Th17 cells as compared to 1C6F Th17 cells is a compelling finding. We find that 18 transcripts are differentially regulated between H_f_XX and 1C6 female Th17 cells. While the modest number of DEG were perhaps surprising, given the stark phenotypic differences elicited by these cells, these genes included several epigenetic modifiers and transcription factors that included Runx2. It had previously been demonstrated that deletion of Runx2 in activated but unskewed CD4^+^ T cells augments their expression of RORγt^95^. Here, we show that enforced expression of Runx2 in differentiated Th17 cells promotes lineage deviation and reduces their pathogenicity in vivo. These findings collectively establish Runx2 as a determinant of Th17 cell fate and encephalitogenic potential.

More fundamental was the question of why H_f_XX and WT female Th17 cells give rise to such different phenotypes despite sharing the same genotype. By globally inhibiting DNA methylation, we elicited an immunomodulatory phenotype in H_f_XX Th17 cells, characterized by specific upregulation of IL-10 and β-catenin. Both H_f_XX and WT female mice have WT female dams; however, H_f_XX mice have H_m_XY^KO^ sires. Our data raise the possibility that heritable epigenetic marks might underpin the unexpectedly severe pathogenicity of H_f_XX Th17 cells. Parent-of-origin imprinting effects have been uncovered in active immunization models of EAE^65,96,97^ and patient cohort studies have indicated that they might underlie a non-Mendelian pattern of MS susceptibility loci^64^. In the future, it will be interesting to further explore the possibility that parental genetic imprinting can influence the function of Th17 cells in autoimmune disease.

In conclusion, using a T cell adoptive transfer model of progressive MS, we demonstrate that Th17 cell pathogenicity is governed by an interplay between hormonal sex, chromosomal sex, and epigenetic regulation. Androgen signaling promotes a mildly pathogenic Th17 phenotype through a Wnt-associated pathway, while specific X-linked factors similarly restrain disease. These findings suggest that the balance between protective and pathogenic sex-dependent programs, rather than biological sex itself, may determine outcomes in autoimmune diseases such progressive MS.

## Acknowledgments

We thank Loïc Mangnier for assistance with bioinformatics; Vincent Desrosiers of the CRCHU de Québec-Université Laval Flow Cytometry Platform for cell sorting assistance; Kim Larose-Labrecque, Mathieu Vallière-St-Amant and Sonia Francoeur for animal care. 1C6 mice were provided by Vijay K. Kuchroo. M.U., I.A., P.M.I.A.D. and V.S. were supported by doctoral studentships from MS Canada. M.R.F. was supported by a doctoral studentship from the Fonds de recherche de Québec-Santé (FRQS). A.F. was supported by a master’s studentship from the FRQS. M.R. was supported by a Senior *«chercheurs-boursiers»* salary support award from the FRQS. The work was funded by an Operating Grant (#3781) from MS Canada, a Project Grant (#PJT191944) from the Canadian Institutes of Health Research and a Catalyst Grant (#1253196) from MS Canada. Biorender was used to generate some schematics.

---

**Supplementary Figure 1.**
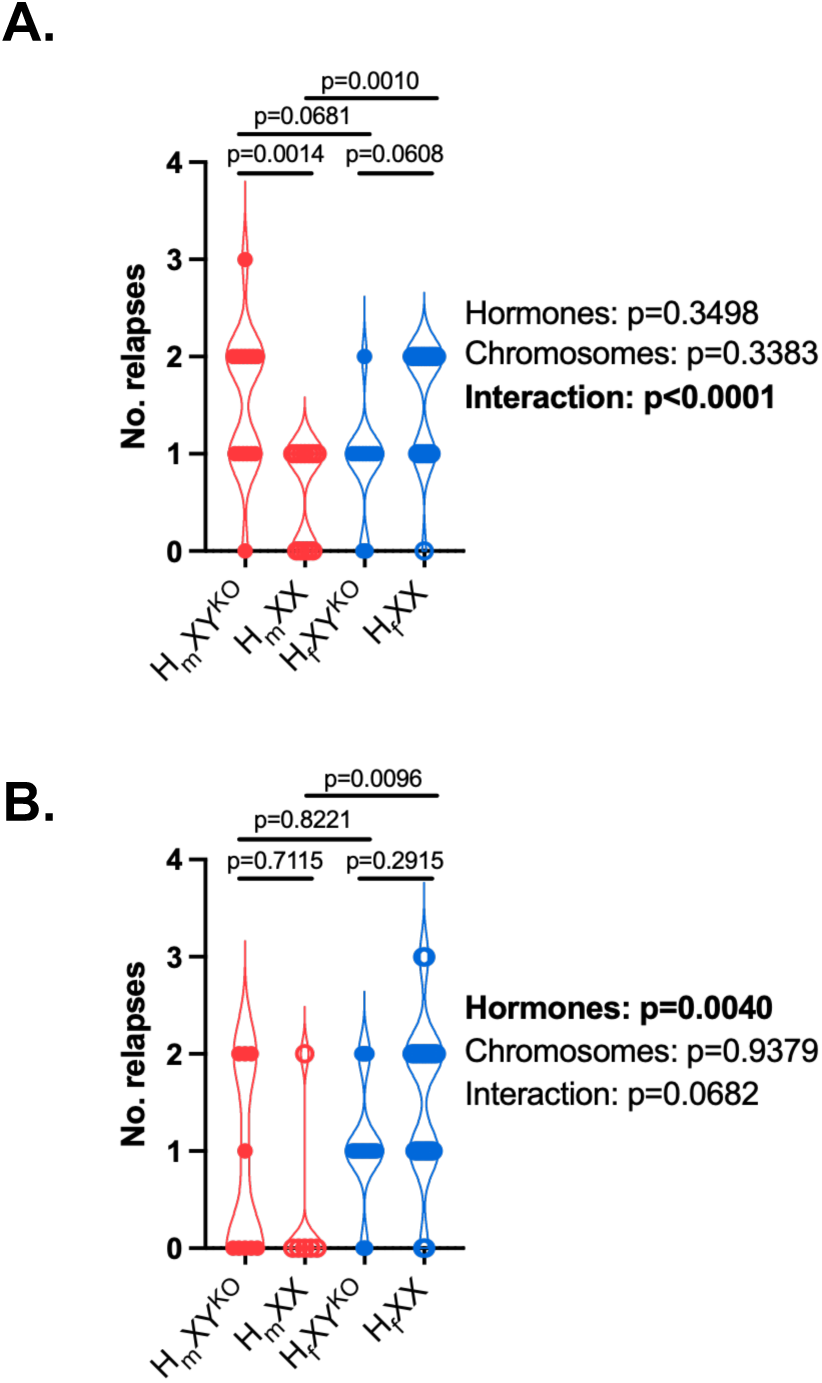
Data linked to Figure 1. **A, B.** Comparison of number of relapses in female (**A**) or male (**B**) recipients of FCG Th17 cells. Hormone, chromosome and interaction effects were measured by two-way ANOVA. Pairwise comparisons were made by Tukey’s post-hoc test. Linked to main Figure 1.

**Supplementary Figure 2.**
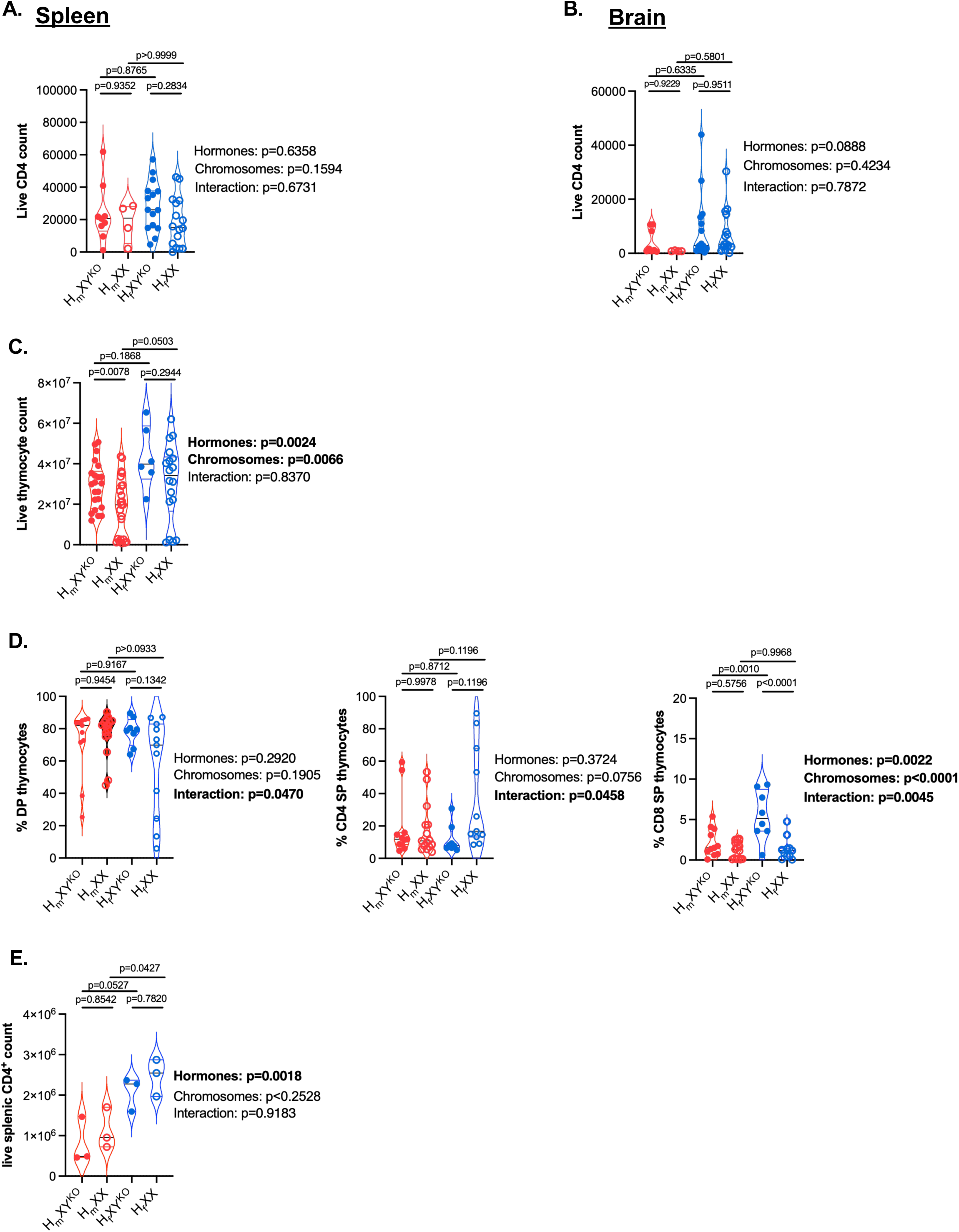
Data linked to Figure 2. **A, B.** Live CD4^+^ count in spleens (**A**) and brain (**B**) of male NOD.*Scid* recipients of FCG:1C6 Th17 cells. H_f_XX, n=16; H_f_XY^KO^, n=15; H_m_XX, n=4; H_m_XY^KO^, n=8. **C.** Live thymocyte counts from FCG:1C6 mice. H_m_XX, n=24; H_m_XY^KO^, n=22; H_f_XX, n=18; H_f_XY^KO^, n=6. **D.** Frequency of DP, CD4^+^ SP and CD8^+^ SP thymocytes from FCG:1C6 mice. H_m_XX, n=15; H_m_XY^KO^, n=11; H_f_XX, n=11; H_f_XY^KO^, n=8. **E.** Absolute count splenic CD4^+^ T cells from FCG:1C6 mice. n=3, each genotype. Hormone, chromosome and interaction effects were measured by two-way ANOVA. Pairwise comparisons were made by Tukey’s post-hoc test. Linked to main Figure 2.

**Supplementary Figure 3.**
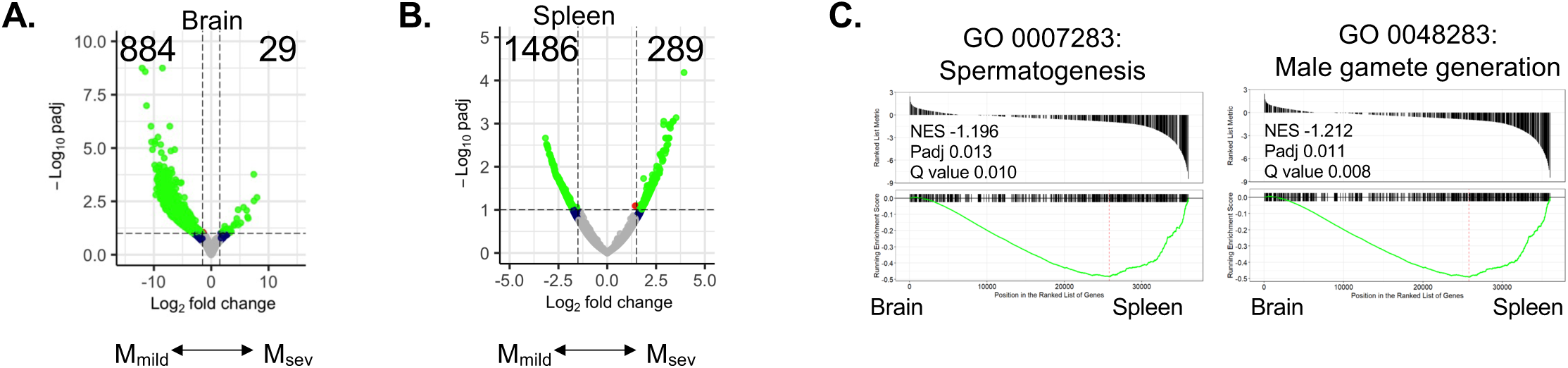
Data linked to Figure 3. **A, B.** Volcano plots comparing M_sev_ and M_mild_ T cell transcriptomes from brain (**A**) and spleen (**B**). P_adj_ and log_2_ fold change cut-off were set at 0.1 and 1.5, respectively. **C.** NES plots for the GO terms “spermatogenesis” and “male gamete generation” upon comparison of pooled M_sev_ and M_mild_ brain vs pooled M_sev_ and M_mild_ splenic T cell transcriptomes.

**Supplementary Figure 4:**
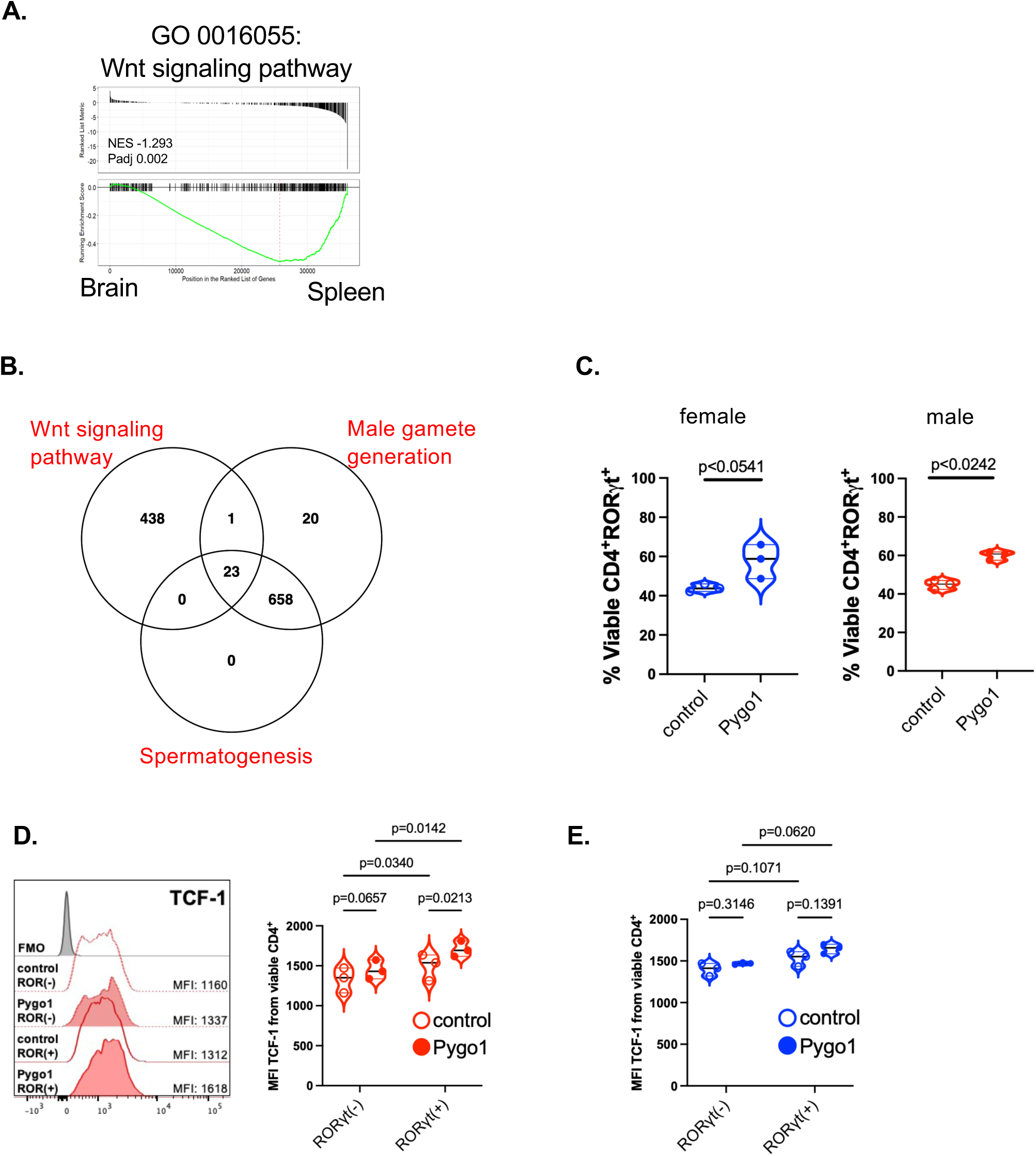
Data linked to Figure 4. **A.** GSEA plots for the GO term “Wnt signaling pathway” comparing all male brain samples (M_sev_ and M_mild_, n=3 each) to all male spleen samples. **B.** Venn diagram displaying common genes tagged in GO terms “spermatogenesis” (GO:0007283), “male gamete generation” (GO:0048232), and “Wnt signaling pathway” (GO:0016055) **C.** Quantification of RORγt expression from female (left) or male (right) 1C6 Th17 cells transduced with control- or Pygo1-overexpressing retrovirus (n=3 each sample) **D, E.** WT male (**D**) or female (**E**) 1C6 Th17 cells were transduced with control- or Pygo1-expressing retrovirus. Representative plots and quantification of TCF-1 expression from RORγt^+^ and RORγt^-^ gated cells after 5 days of culture (n=3 each condition). Quantifications based on FACS gates set on viable CD4^+^ events. **C**, paired *t*-test. **D, E**. Sidak’s multiple comparisons test.

**Supplementary Figure 5:**
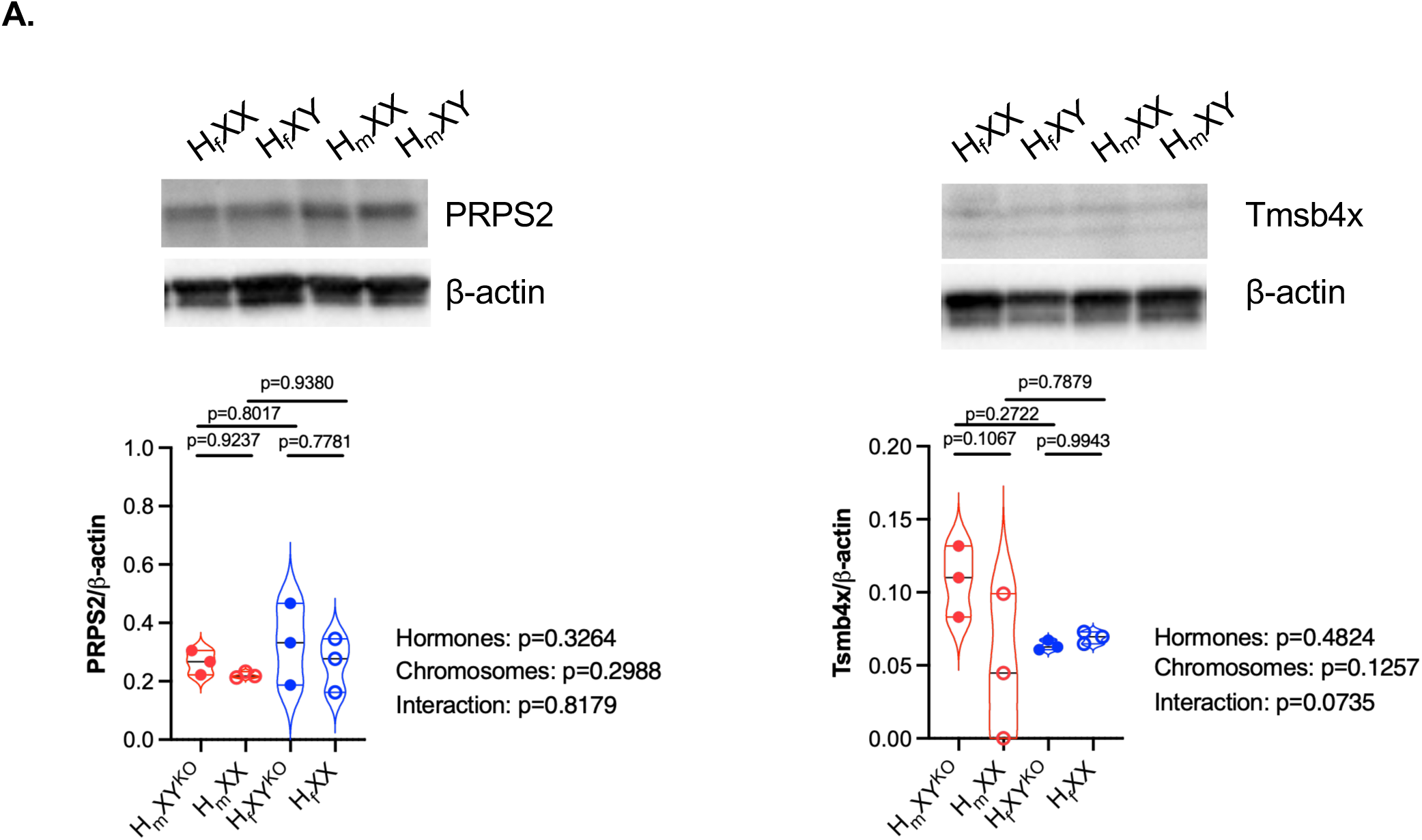
Data linked to Figure 5. **A** Protein expression of PRPS2 and TMSB4X (**C**) from FCG Th17 cells (n=3, each genotype). Representative images from one mouse of each genotype. Quantification of image density normalized to beta-actin. Each symbol represents a culture derived from an individual mouse. Hormone, chromosome and interaction effects were measured by two-way ANOVA while pairwise comparisons were made by Tukey’s post-hoc test.

**Supplementary Figure 6:**
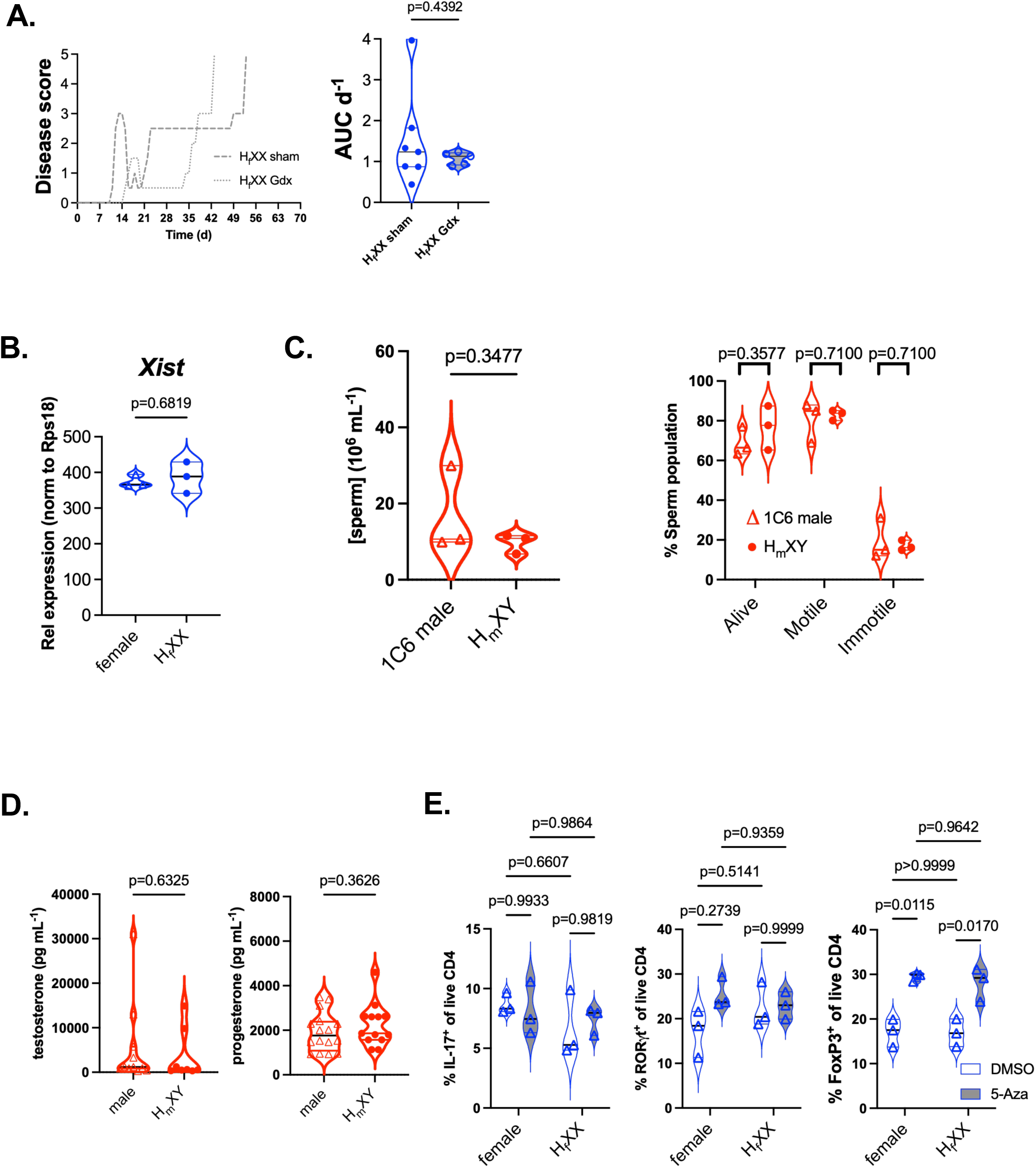
Data linked to Figure 6. **A.** H_f_XX:1C6 mice were gonadectomized or sham-operated, and Th17 cells were derived from these mice and adoptively transferred to female NOD.Scid recipients. Disease burden and rapidity were quantified. n=7, recipients of sham-operated H_f_XX 1C6; n=5, recipients of Gdx-H_f_XX 1C6. Data from a single transfer. **B.** qPCR of *Xist* expression from H_f_XX and 1C6F Th17 cells (n=3 each). **C.** Quantification of sperm concentration, live percentage and motility from 1C6 males and H_m_XY (n=3 each). Data analyzed by unpaired *t*-test. **D.** Serum hormone concentration of testosterone and progesterone from 1C6 males (n=11) and H_m_XY (n=8). Data analyzed by unpaired *t*-test. **E.** Quantification of expression of IL-17, RORγt and FoxP3 from WT female 1C6 and H_f_XX 1C6 cultures studied in Figure 6F. Each symbol represents an individual mouse. **A-D**, unpaired two-tailed *t*-test. **E,** Sidak’s multiple comparisons test.

## Materials and methods

### Mice

1C6 mice have been previously described^18^. FCG mice (B6.Cg-Tg(Sry)2Ei *Sry^dl1Rlb^* T(XTmsb4x-Hccs;Y)1Dto/ArnoJ; strain #010905) were obtained from Jackson Labs and were backcrossed to the NOD genetic background for 5 generations using Marker Assisted Accelerated Backcrossing (MAX-BAX; Charles River Laboratories). A 348-marker SNP panel revealed a 99.5% match between backcrossed FCG mice and WT NOD male mice. FCG NOD mice were then crossed to the 1C6 line on the NOD background to generate FCG:1C6 mice. NOD.*Scid* mice were purchased from Jackson Labs and were bred in our animal facility. All mice were housed in the animal facility of the Centre de recherche du CHU de Québec-Université Laval on a 12-hour light dark cycle and were supplied with ad libitum food and water. NOD*.Scid* mice received sterile water supplemented with antibiotics (trimethoprim 0.4 mg ml^-1^ + sulfamethoxazole 0.08 mg ml^-1^) along with sterile food. When required, gonadectomy procedures were performed by trained animal staff on 21-24-day old mice and sperm isolation was performed on 8-12 week old mice. All animal studies were reviewed and approved by the relevant Animal Care Committee (CPAUL-3) of Université Laval.

### Th17 cell differentiation and culture

Naïve CD4 cells were enriched from single cell suspensions of lymph nodes and spleen of 8-12-week-old mice using anti-CD4 Microbeads (Miltenyi Biotec) followed by high-speed cell sorting (FACSAria II; BD Biosciences) of live CD4^+^CD62L^+^ T cells. For plate-bound conditions, culture plates were incubated overnight with 2 µg ml^-1^ each of anti-CD3 (clone 1452c11, BioXcell) and anti-CD28 (clone 37.51, BioXcell) antibodies in PBS (Corning) at 4°C. For MLR or same-sex APC stimulation, APCs were first generated by irradiating (2000 cGy) splenic and lymph node CD4^neg^ cells from splenocytes and lymph nodes. Irradiated APCs were then co-cultured with naïve CD4^+^ T cells at a ratio of 5:1 with soluble CD3 (2 µg ml^-1^). Plate-bound and APC-stimulated naïve CD4^+^ T cells were differentiated under Th17 conditions as previously described (Doss et al., 2021): 20 ng mL^-1^ rmIL-6 (Miltenyi Biotec), 3 ng mL^-1^ rhTGFβ (Miltenyi Biotec) and 20 µg ml^-1^ anti-IFNγ (clone XMG1.2 BioXcell). On day 2, cells were transferred to an uncoated culture plate in the presence of 20 ng mL-1 rmIL-23 (R&D Systems) and then cultured for an additional 3 days (5 days total). Testosterone, DHT and estradiol (E2) were kind gifts of Dr. Donald Poirier (Université Laval). F5446 was purchased from MedChemExpress and 5-azacytidine was purchased from STEMCELL Technologies. These reagents were applied at d0 of Th17 culture, at the indicated concentrations, and were maintained for 5 days’ culture.

### Retrovirally-mediated gene transfer

For overexpression studies, Pygo1 or Runx2 cDNA were cloned into the pMSCV-IRES-GFP (pMIG) retroviral expression vector that harbors a bicistronic sequence allowing the co-expression of enhanced green fluorescent protein (eGFP) as a reporter. For knockdown studies, a sequence targeting exon 3 of Suv39h1 (5′-CTCTGCATCTTCCGCACTAAT-3′) was identified using the Genetic Perturbation Platform (GPP) Web Portal. An shRNA oligonucleotide with *EcoRI*- and *XhoI*-compatible overhangs (IDT) was designed in a miR-30 backbone format^98^, in which the target sequence is embedded within flanking sequences that mimic the endogenous miR-30 hairpin structure, and was ligated upstream of the eGFP reporter in the pMSCV-IRES-GFP II (pMIG II) retroviral vector backbone (Addgene plasmid #52107).We followed our previously described transduction protocol^14,99^. Briefly, RV particles were produced by transfecting (XtremeGENE HP, Sigma) HEK293T cells (ATCC) with cDNA-encoding pMIG or shRNA-encoding LMP along with EcoPCL and gag-pol. RV particle-containing supernatant were layered on Th17 cells at d2 of T-cell culture in the presence of polybrene (8 μg mL^-1^, Sigma) and were spin-infected (2000g, 90’, 32°C). Bicistronic reporter expression was assessed at d5 of T cell culture by flow cytometric analysis or high-speed cell sorting.

### EAE induction, scoring and quantification

NOD.*Scid* recipient mice (10-12 weeks old) received, i.v., 5 × 10^6^ Th17 cells generated under plate-bound conditions. To induce EAE from RV-transduced Th17 cells, we sorted bicistronic reporter-positive CD4^+^ Th17 cells after 5 days’ culture and injected 1.5 × 10^6^ cells to NOD.*Scid* mice. In all EAE experiments, recipient mice received 200 ng of pertussis toxin (List Biological Labs) i.p. on day 0 and day 2. Disease score and relapse/remission criteria were assessed daily as previously described^14^. Briefly, for classical disease: score 0, no disease; score 1, lack of tail tonicity; score 2, lack of righting reflex; score 3, hind limb paralysis; score 4, front limb paralysis; score 5, ethical endpoints attained. Mice with ataxic/atypical symptoms were scored as previously reported^100^: score 0, no disease; score 1, slight head tilt; score 2, pronounced head tilt; score 3; inability to maintain a straight walking trajectory; score 4, lateral recumbency; score 4.5, continuous rolling; score 5, ethical endpoints attained. Mice were weighed at the beginning of the protocol and then daily upon the start of disease symptoms. Mice with advanced symptoms were discussed daily with the veterinary service of Université Laval and were humanely euthanized if it was deemed that ethical endpoints had been reached. “Disease onset” was defined as the protocol day on which the first sign of disease was observed. “Relapse” was defined as an increase in clinical score of ≥1 lasting at least two days, excluding the initial attack; “remission” was defined as a decrease in score of ≥1 lasting at least two days^14^. Disease burden (AUC d^-1^) was calculated by first assessing the AUC of individual disease curves until the day of sacrifice (either day that ethical endpoints were attained, or experimental endpoint of d70). AUC d^-1^ was then determined by using the day of sacrifice as the denominator. Rapidity was assessed by determining the slope of the linear regression line for each disease curve, from d0 of the protocol to the day of sacrifice. For ex vivo mRNA-seq analyses, M_sev_ mice were sacrificed as they attained ethical endpoints on d35 (n=1) and d58 (n=2) of EAE protocol. M_mild_ and F mice had disease severity of 0.5-2 and were sacrificed in parallel to M_sev_ mice.

### Mononuclear cell isolation

Mice were euthanized and perfused with cold PBS. Spleens and thymi were mashed through a 70 μm strainer. Splenic single cell suspensions were treated with ACK buffer (Gibco) to lyse red blood cells. Brains was isolated from dissected skulls, and spinal cords were flushed from vertebral columns using cold PBS. Tissues were mashed mechanically using a PTFE tissue grinder (VWR) and homogenization solution (4ng ml^-1^ liberase + and 25 ng ml^-1^ DNase in HBSS) was used for tissue digestion. After 30 mins, the mixture was passed through a 70µM cell strainer. The filtrate was further resuspended in 35% Percoll and subjected to centrifugation at 2500 rpm for 10 minutes at RT. The isolated mononuclear cells were then washed with PBS and subsequently used for downstream applications.

### Flow cytometry

Antibodies to surface markers (CD4 [clone RM4-5, Thermo Fisher Scientific], CD8 [clone 53-6.7, Thermo Fisher Scientific], CD25 [clone PC61, BioLegend], Vβ7 [clone TR310, BioLegend], CD44 [clone IM7, BioLegend], CD62L [clone MEL-14, Invitrogen], Slamf6 [clone 13G3-19D, Invitrogen], CXCR6 [clone DANID2, Invitrogen]) and intracellular markers (GFP [clone FM264G, BioLegend], IL-17 [clone TC11-18H10.1, bioLegend], IFNγ [clone XMG1.2, Thermo Fisher Scientific], GM-CSF [clone MP1-22E9, Thermo Fisher Scientific], IL-2 [clone JES65114, BioLegend], TNFα [clone MP6-XT22, Thermo Fisher Scientific], IL-10 [clone JES5-16E3, BioLegend], RORγt [clone AFKS-9, Thermo Fisher Scientific], T-bet [clone 4B10, BioLegend], TCF-1 [clone C63D9, Cell Signaling], FOXP3 [clone NRRF-30, Invitrogen], β-catenin [clone 1588, Thermo Fisher Scientific], AR [clone 523339, R&D Biosystems], ER1 [clone E115, Abcam], Runx2 [clone EPR14334, Abcam]) were used. For surface markers, cells were briefly incubated with FC Block (BioLegend) prior to staining with fluorescently antibodies (30’). Cells were subsequently washed and stained with Fixable Viability Dye (ThermoFisher Scientific). For intracellular (ICC) staining, cells were first pretreated with phorbol-12-myristate-13-acetate (PMA; Sigma, 50 ng mL^-1^), ionomycin (Sigma, 1µM) and Golgi stop (BD Biosciences,1 µL per mL of media) for 4 hours, prior to being washed twice with PBS. After Fc Block and staining with surface antibodies and Fixable Viability as above, cells were washed and fixed in the appropriate fixation buffer. Subsequent stainings were conducted in the appropriate perm/wash buffer. For cytokines, Fixation Buffer (cat 420801; BioLegend) and Perm Wash Buffer(Cat 421002; BioLegend) were used; for AR, ER, RORγt, Tbet and Runx2, True-Nuclear Transcription Factor Buffer Set (Cat: 424401, BioLegend) was used; and for β-catenin and FoxP3, a FOXP3/Perm Buffer Set (Cat: 421403, BioLegend) were used. Absolute cell count was measured by using 123count ebeads (Thermo Fisher Scientific). Samples were processed on LSRII or FACSCanto flow cytometers (BD Biosciences) or on a FACSAria Fusion cell sorter (BD Biosciences) Data was analyzed using Flowjo (BD Biosciences) and gates were based on florescence minus one (FMO) controls.

### mRNA-seq

For mRNA-seq on ex vivo cells, splenic and brain mononuclear cells were isolated as described above and live CD4^+^ cells were purified by high-speed cell sorting. After extraction (TRIzol, Thermo Fisher Scientific) RNA was subjected to the NEBNext Ultra II RNA Library Prep Kit (New England Biolabs) for library preparation. For mRNA-seq on in vitro-cultured cells, Th17 cells from day 5 of the culture were washed, and RNA was isolated using the RNeasy Kit (Qiagen). NEBNext Ultra II directional RNA Library Prep Kit for Illumina (New Englands Biolabs Inc., Ipswich, MA, USA) was then used to prepare mRNA sequencing libraries. Briefly, 1 ng of total RNA were purified using the NEBNext poly(A) mRNA Magnetic Isolation module (New Englands Biolabs Inc., Ipswich, MA, USA) and used as a template for cDNA synthesis by reverse transcriptase with random primers. The specificity of the strand was obtained by replacing the dTTP with the dUTP. The quality of final amplified libraries was examined with a DNA ScreenTape D1000 assay on a TapeStation 2200 (both Agilent). Libraries were then subjected to paired-end 100 pb sequencing on an Illumina NovaSeq 6000 at a depth of 20 million (ex vivo Th17) or 25 million (in vitro Th17) reads at the Next-Generation Sequencing Platform, Genomics Center, CHU de Québec-Université Laval Research Center. Each experimental group contained a minimum of three biological replicates.

### Bioinformatics analysis

We used trimmomatic v0.39 for trimming fastq files with the following command: java -jar trimmomatic-0.39.jar PE -phred 33 sample_R1.fastq sample_R2.fastq trimmed/paired_sample_R1.fastq trimmed/unpaired_sample_R1.fastq trimmed/paired_sample_R2.fastq trimmed/unpaired_sample_R2.fastq ILLUMINACLIP:TruSeq3-PE.fa:2:30:10\LEADING:26\ TRAILING:26\ MINLEN:30

The trimmed FASTQ files were aligned to the Mus musculus reference genome (Ensembl; Mus.musculus.GRCm39.cds.all.fa.idx) and quantified using Kallisto version 0.44.0. Differential analysis was conducted in RStudio using the DESeq2 package to assess mRNA gene expression differences across various groups. The annotation of the ENSEMBLE IDs to their respective gene symbols was performed using the “org.Mm.eg.db” package. Additionally, the annotation of the identified genes to their respective chromosomes was conducted with the “TxDb.Mmusculus.UCSC.mm10.knownGene” package. Gene set enrichment analysis (GSEA) was performed using the ‘clusterProfiler’ package in RStudio, with the Benjamini-Hochberg method used to calculate the adjusted p-values (Padj). For data visualization, volcano plots were generated using the ‘EnhancedVolcano’ package in RStudio. Venn diagrams were created using the online tool Venny 2.1.

Further information regarding code related to this study are stored at https://github.com/UmairUHN/Th17-cells-sex-difference. Raw sequencing files are located at Gene Expression Omnibus (series GSE268964, GSE344686).

### Western blotting

Whole cell lysates were generated from FCG Th17 cells with cell lysis buffer (RIPA buffer [Boston Bioproducts], 10% SDS, 100 mM Na3VO4, 100 mM PMSF, 1X Halt Protease/Phosphatase Inhibitor cocktail [Thermo Fisher Scientific]). Clarified and boiled lysates were quantified (DC Protein Assay, Bio-Rad) prior to loading (10 μg/well) on Criterion TGX PreCast gels (Bio-Rad) and transferring to PVDF membrane (Bio-Rad). Membranes were incubated in 5% milk-TBS + 0.05% Tween-20 (60’) to block non-specific binding. Membranes were probed (overnight, 4°C) with rabbit anti-PRPS2 (1:1000, Proteintech) or rabbit anti-Tmsb4x (Invitrogen) followed by HRP-conjugated mouse anti-rabbit (1:10000, Jackson Immunoresearch, 60’). Blots were washed and revealed using Immobilon Western chemilumiscent HRP substrate (Millipore Sigma). Molecular imager ChemiDoc XRS+ (BioRad) was used to detect signal and data were quantified using ImageJ (NIH).

### Quantitative PCR (qPCR)

qPCR for *Xist* performed following our previously established protocol^14^. Briefly, RNA was isolated from Th17 cells using the Direct-zol RNA Purification Kit (Zymo Research), and cDNA was synthesized using the iScript cDNA Synthesis Kit (Bio-Rad). The qPCR was conducted on a LightCycler 480II instrument using TaqMan Gene Expression Master Mix (ThermoFisher Scientific). *Rps18* was utilized as a reference control.

### Sex hormone analysis from serum

Serum was isolated from the blood of mice aged 8-12 weeks and processed for liquid chromatography with tandem mass spectrometry (LC-MS/MS) at Pharmacogenomics Laboratory of Centre de recherche du CHU de Québec-Université Laval. Sex hormones were assessed following the previously described approaches of Tiwari et al.^101^ and Audet-Delage et al.^102^ Further details are in the Supplementary Methods.

### Data availability

Raw data are available upon reasonable request to the corresponding author. Sequencing data (GEO) will be made public upon manuscript acceptance.

