## Supplementary Table 1 for "The antagonism of sex chromosomes and sex hormones dictates outcomes in Th17 cell-mediated central nervous system autoimmunity"

| **Gene** | **Fold change** | **P_adj_** | **Mean read count** |
| --- | --- | --- | --- |
| *Prps2* | 2.372 | 1.48 x 10^-21^ | 773.55 |
| *Tmsb4x* | 2.040 | 2.89 x 10^-12^ | 24700.54 |
| *Msl3* | 1.866 | 1.17 x 10^-8^ | 1010.42 |
| *Hccs* | 1.727 | 0.001644 | 330.70 |
| *Tlr7* | 1.006 | 0.999804 | 32.72 |
| *Tlr8* | 1 | n/a | 0.16 |
| *Arhgap6* | 1 | 0.999804 | 5.88 |
| *Amelx* | 1 | n/a | 0 |
| *Frmpd4* | 0.999 | 0.999804 | 15.69 |

**Supplementary Table 1. Differential expression of genes within the X-Y chromosomal translocation in FCG Th17 cells.** RNA-seq analysis of the nine genes within the X-Y translocation reported in Pantem et al (2024), comparing chromosomally male (XY) and female (XX) FCG Th17 cells. Genes are ranked by fold change (XY relative to XX), with corresponding adjusted P values and mean normalized read counts.
