## Supplementary Methods for "The antagonism of sex chromosomes and sex hormones dictates outcomes in Th17 cell-mediated central nervous system autoimmunity"

- Serum analysis of sex hormones
- Sperm motility and viability assessment

**Serum analysis of sex hormones**

Briefly, serum samples were mixed 0.5M NH_4_CH_3_CO_2_, internal standard (Estrone-d4, Estradiol-d4, DHT-d3, PROG-d9, A4-d7 and Testosterone-d3) and 1-chlorobutane prior to vortexing. After centrifugation, the organic layer containing steroids was transferred and evaporated under nitrogen at 45 °C for 10 min with a Turbo Vap (Zymark Corporation, Hopkinton, MA, USA). To enhance detection of estrogens, a derivatization procedure was applied using dansyl chloride (0.5 mg/mL final in 50% acetone and 50 mM sodium bicarbonate, pH 9.0), as described previously^100^. Samples were heated for 5 min at 60 °C, mixed with water:methanol (80:20, v/v) and loaded on Strata X 60 mg columns (Phenomenex, Torrance, CA, USA). After being washed with water and water:methanol (45:55, v/v), PROG and androgens were eluted with water:methanol (10:90, v/v), evaporated to dryness at 35°C under nitrogen gas, reconstituted in 100 uL of water: methanol, (50:50, v/v) and injected into the LC-MS/MS system. The estrogens were eluted with dichloromethane:methanol (50:50, v/v). Eluates were evaporated to dryness at 45 °C under nitrogen gas, reconstituted in 100 μL of acetone:water (75:25, v/v) and injected into a LC-MS/MS system. Analytical procedures were performed with a HPLC Nexera system (Shimadzu Scientific Instruments, Columbia, MD, USA) and a Sciex mass spectrometer 6500 triple quadrupole system (Concord, ON, Canada) equipped with a turbo ion-spray source, configured in the multiple reaction monitoring (MRM) mode. The chromatographic separation was achieved as described for androgens and progesterone^100^ with some modifications. Estrogen separation was achieved as described^101^ with some modifications. Separation was performed on a Supelco Ascentis Express C18 column (150 ×3.0 mm ID, particle size 2.7 μm; Sigma, Oakville, ON, Canada). A binary mobile phase, consisting of 0.1% formic acid in H2O (solvent A), and 0.1% formic acid in ACN (solvent B), was used at a flow rate of 0.8mL/min. The analytes were eluted using the following program: 0–23 min, linear gradient 40–80% B; 23.0-23.1 min, linear gradient 80–95%; 23.1-27.0 min, isocratic 95% B; 27.0–27.1 min, linear gradient 95–40% B; 27.1–34 min, isocratic 40% B to re-equilibrate the column. The transitions were 291 🡪 255 m/z for DHT, 315.1 🡪 97.1 m/z for progesterone, 287 🡪 97 m/z for A4, 504 🡪 171 m/z for estrone and 506 🡪 171 m/z for estradiol.

**Sperm motility and viability assessment**

Sperm motility was evaluated using a computer-assisted semen analysis system (CASA; CEROS version 12, Hamilton-Thorne Research Inc., Beverly, MA, USA). Cauda epididymal spermatozoa were released into pre-warmed Whitten's medium and maintained at 37°C prior to analysis. Sperm concentration was determined, and motility was assessed at 37°C using 20-μm-deep analysis chambers (Cell Vision, CV1020-2CV, Netherlands). CASA settings were configured according to the CEROS technical guide version 12 for mouse spermatozoa. For each sample, a minimum of 200 spermatozoa distributed across at least five randomly selected fields were analyzed. Motility parameters recorded included total motility (%) and progressive motility (%). Sperm viability was assessed by staining spermatozoa with SYBR-14 (100 nM) for 10 min at 37°C, followed by propidium iodide (PI; 12 μM) for an additional 5 min in the dark. Samples were analyzed by fluorescence microscopy using the CASA system. Live spermatozoa were identified by green SYBR-14 fluorescence, whereas non-viable spermatozoa were identified by red PI fluorescence. At least 200 spermatozoa were counted per sample, and viability was expressed as the percentage of live sperm relative to the total number of sperm analyzed.
